# Hippocampal sequences span experience relative to rewards

**DOI:** 10.1101/2023.12.27.573490

**Authors:** Marielena Sosa, Mark H. Plitt, Lisa M. Giocomo

**Author notes:** Co-corresponding authors: Marielena Sosa; Lisa Giocomo.

## Abstract

Hippocampal place cells fire in sequences that span spatial environments and non-spatial modalities, suggesting that hippocampal activity can anchor to the most behaviorally salient aspects of experience. As reward is a highly salient event, we hypothesized that sequences of hippocampal activity can anchor to rewards. To test this, we performed two-photon imaging of hippocampal CA1 neurons as mice navigated virtual environments with changing hidden reward locations. When the reward moved, the firing fields of a subpopulation of cells moved to the same relative position with respect to reward, constructing a sequence of reward-relative cells that spanned the entire task structure. The density of these reward-relative sequences increased with task experience as additional neurons were recruited to the reward-relative population. Conversely, a largely separate subpopulation maintained a spatially-based place code. These findings thus reveal separate hippocampal ensembles can flexibly encode multiple behaviorally salient reference frames, reflecting the structure of the experience.

## INTRODUCTION

Memories of positive experiences are essential for reinforcing rewarding behaviors. These memories must also have the capacity to update when knowledge of reward changes (i.e. a water source moves to a new location)^1,2^. How does the brain both amplify memories of events surrounding a reward and maintain a stable representation of the external world? The hippocampus provides a potential neural circuit for this process. Hippocampal place cells fire in one or a few spatially restricted locations, defined as their place fields^3^. In different environments, place cells “remap”, such that the preferred location or firing rate of their place field changes^4–12^. Together, the population of place cells creates reliable sequences of activity as an animal traverses an environment, resulting in a unique neural code for each environment and a unique sequence of spiking for each traversal^9,13–21^. While these patterns support spatial navigation^22–26^, recent evidence has revealed that hippocampal neurons also show reliable tuning relative to a variety of non-spatial modalities and create sequences of activity for these modalities^17,27,28^. For instance, sequential hippocampal firing has been observed across time^29–35^, in relation to progressions of auditory^36^, olfactory^37–42^, or multisensory stimuli^43–45^, and during accumulation of evidence for decision-making tasks^46^. Together, these findings suggest that the hippocampus can be understood as generating sequential activity^13,17,27,29,47,48^ to encode the progression of events as they unfold in a given experience^27,28,49–51^.

Multiple lines of evidence demonstrate that the hippocampus prioritizes coding for aspects of experience that are particularly salient or that can affect the animal’s behavior^27,33–46,52–62^. The presence of food or water reward is one such highly salient event that is consistently prioritized, as demonstrated by prior work finding that place cells tend to cluster near (i.e. “over-represent”) reward locations^1,2,54,55,63–66^. In addition, a small subpopulation of hippocampal cells are active precisely at rewarded locations^54,64^, even when the reward is moved to a new location^54^. Moreover, optogenetic activation of cells with place fields near reward drives the animal to engage in reward-seeking actions^24^, suggesting a causal role for hippocampal activity in reward-related behaviors. Yet it remains unclear whether the hippocampus encodes complete sequences of events around rewards separately from the influence of other sensory stimuli. Moreover, it is incompletely understood how different aspects of experience are represented simultaneously.

We therefore hypothesized that reward may anchor the sequential activity of the hippocampus in a subpopulation of pyramidal cells. Remapping dynamics supporting this hypothesis have been reported at locations very close to reward sites^1,2,54,55,63–66^, but in order to support memory of events leading up to and following rewarding experiences, hippocampal activity encoding events at distances further away from reward must also be able to update when reward conditions change. Yet predictable remapping that spans the entire environment in response to changing reward locations has not been demonstrated. We reasoned that previously reported reward-specific cells^54^ may comprise a subset of a larger population encoding an entire sequence of events relative to reward. This hypothesis predicts that moving the reward within a constant environment should induce remapping even at locations far from reward, in a manner that preserves sequential firing relative to each reward location. In parallel, we predicted that a subset of hippocampal neurons should preserve their firing relative to the spatial environment, reflecting an ability of the hippocampus to flexibly anchor to both the spatial environment and the experience of finding reward as two salient reference frames^45,59,60,67–73^ that are required to solve the task at hand.

To investigate whether the hippocampus simultaneously encodes sequential experience relative to reward and the spatial environment, we used 2-photon (2P) calcium imaging^74,75^ to monitor large populations of CA1 neurons in head-fixed mice learning a virtual reality (VR) task. The use of VR provided tight control of the sensory stimuli and constrained the animal’s trajectory, allowing us to more readily observe remapping of neurons in relation to distant rewards, a phenomenon potentially less visible in freely moving scenarios. Further, the task included multiple updates to hidden reward zones across two environments, allowing us to dissociate spatially-driven and reward-driven remapping. We found a subpopulation of neurons that, when rewards moved, remapped to the same relative position with respect to reward and thus constructed a sequence of activity relative to the reward location. With increasing experience, more neurons were recruited to these “reward-relative sequences”. These results suggest that the hippocampus learned a generalized representation of the task anchored to reward, while also maintaining a spatial map in largely separate neural ensembles.

## RESULTS

### Monitoring neural activity in mice during a virtual reality navigation and reward learning task

We performed 2P calcium imaging to monitor CA1 neurons expressing GCaMP7f in head-fixed mice learning a VR linear navigation task (**Fig. 1A-C**). To observe putative remapping related to rewards, and examine how this remapping might develop with experience, we designed a task with multiple changing hidden reward locations. In the task, animals were required to run down a 450 cm virtual linear environment (Env 1) with a single hidden, 50-cm reward zone. Sucrose water reward was delivered operantly for licking within the hidden reward zone (**Fig. 1D-E**). Imaging began on day 1 of task acquisition. On day 3, the reward zone was moved within a “switch” session to a new location on the track. The reward switch was signaled by automatic reward delivery on the first 10 trials after the switch only if the mouse failed to lick in the new zone. On day 5, a third reward zone was introduced, followed by a return to the original zone on day 7. On day 8, the reward switch coincided with the introduction of a novel environment (Env 2) that evokes spatially-driven global remapping^7^. The sequence of switches was subsequently reversed in Env 2 (**Fig. 1E**). On intervening non-switch (“stay”) days, the last reward location from the previous day was maintained. Reward switch sequences were counterbalanced across animals (n=7 “switch” mice, **Fig. S1A-D**). The three possible reward locations were positioned equidistant from each of the “tower” landmarks along the track, thus controlling for proximity of each reward to visual landmarks^8,55,76–80^. A separate “fixed-condition” group of mice (n=3) experienced only one reward location in a single environment (Env 1) (**Fig. S1D, F**), to control for the effect of extended experience alone compared to learning about the reward switches. To keep animals engaged and prevent the movement of the reward zone from being a completely novel experience, on ∼15% of trials the reward was randomly omitted for all animals. At the end of the track, the mouse passed through a brief gray “teleport” tunnel (∼50 cm + temporal jitter up to 10 s; see Methods) before starting the next lap in the same direction. Time in the teleport tunnel was randomly jittered so that exact time of track entry could not be anticipated.

**Figure 1.**
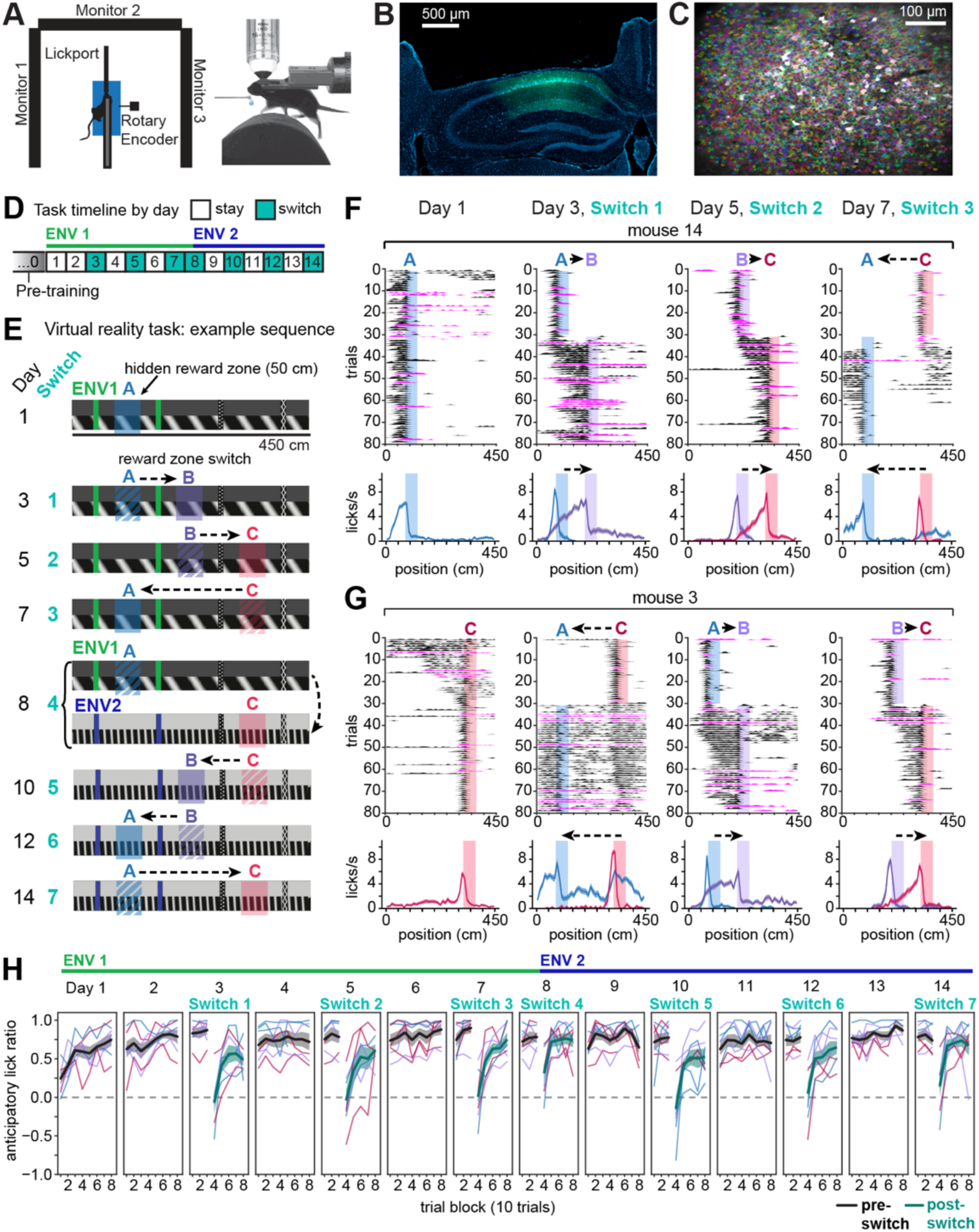
2P imaging of CA1 neurons in a VR task with changing hidden reward locations. **A)** Top-down view (left) and side view (right) of head-fixed VR set-up. 5% sucrose water reward is automatically delivered when the mouse licks a capacitive lick port. **B)** Coronal histology showing imaging cannula implantation site and calcium indicator GCaMP7f expression under the pan-neuronal synapsin promoter (AAV1-Syn-jGCaMP7f, green) in dorsal CA1 (DAPI, blue). **C)** Example field of view (mean image) from the same mouse in (B) (mouse m12) with identified CA1 neurons in shaded colors (n=1780 cells). **D)** Task timeline. On “stay” days, the reward remains at the same location throughout the session; on “switch” days, it moves to a new location after the first 30 trials (typically 80 trials/session). **E)** Side views of virtual linear tracks showing an example reward switch sequence. Shaded regions indicate hidden reward zones, for illustration only. On intervening days, the last reward location from the previous day is maintained. **F)** Licking behavior for an example mouse (m14) on the first day and first 3 switches. Top row: Smoothed lick rasters. Rewarded trials in black; omission trials in magenta. Shaded regions indicate the active reward zone. Bottom row: Mean ± s.e.m. lick rate for trials at each reward location (blue=A, purple=B, red=C). **G)** Licking behavior for a second example mouse (m3) that experienced a different starting reward zone and switch sequence from the mouse in (F). See Fig. S1D for the full set of sequence allocations per mouse. **H)** Behavior across animals performing reward switches (n=7 mice). An anticipatory lick ratio of 1 indicates licking exclusively in the 50 cm before a reward zone (“anticipatory zone”, see Fig. S1E); 0 (horizontal dashed line) indicates chance licking across the whole track outside of the reward zone. Thin lines represent individual mice, with lines colored according to the reward zone active for each mouse on a given set of trials (blue=A, purple=B, red=C); thick black and teal lines show mean ± s.e.m. across mice for pre- and post-switch trials, respectively. See also Figure S1.

We observed that mice developed an anticipatory ramp of licking that, on average, peaked at the beginning of the reward zone (**Fig. 1F-G, Fig. S1A-C**). This is the earliest location where operant licking triggered reward delivery, demonstrating successful task acquisition. After a reward switch, mice entered an exploratory period of licking but then adapted their licking to anticipate each new zone, typically within a session (**Fig. 1F-G, Fig. S1A-C**). We quantified this improvement in licking precision across the switch as the ratio of anticipatory lick rate 50 cm before the currently rewarded zone compared to everywhere else outside the zone (**Fig. S1E**). We found that all mice were able to improve and retain their licking precision for the new reward zone (**Fig. 1H**, n=7 mice), demonstrating accurate spatial learning and memory.

### Moving the reward induces remapping that spans a constant environment

We focused on the switch days to analyze single-cell remapping patterns. First, we identified place cells active either before or after the reward switch (see Methods; means ± standard deviation [s.d.] across 7 mice and 7 switch sessions: 408 ± 216 place cells out of 868 ± 356 cells imaged; 48.6% ± 16.7% place cells active before or after the switch). After a reward switch within an environment, a subset of place cells maintained a stable field at the same location on the track (“track-relative” cells, 20.2% ± 8.8%, mean ± s.d. of all place cells, averaged across all 6 switches within environment, 7 mice; **Fig. 2A**, **Fig. S2A**). However, many place cells remapped, despite the constant visual stimuli and visual landmarks (**Fig. 2A, Fig. S2A-D).** First, we observed subsets of cells with place fields that disappeared (“disappearing”, 11.5% ± 7.8%) or appeared (“appearing”, 8.3% ± 5.2%) after the reward location switched (**Fig. 2A, Fig. S2A**). Other cells precisely followed the reward location (“remap near reward”, 4.7% ± 3.9%, **Fig. 2A, Fig. S2A- D**), consistently firing within ±50 cm of the beginning of the reward zone, similar to previous reports^54^. Notably, a subset of place cells with fields distant from reward (>50 cm) also remapped after the reward switch (“remap far from reward”, 15.6% ± 3.7%; **Fig. 2A, Fig. S2A**, right). A remaining 39.9% ± 19.3% of place cells did not show sufficiently stereotyped remapping patterns to be classified by these criteria. At the population level, we observed that a reward switch within a constant environment induced more remapping than the spontaneous instability seen with a fixed reward and fixed environment^81–85^, but less remapping than the introduction of a novel environment with the reward switch^5–11^ (**Fig. S2B-D**).

**Figure 2.**
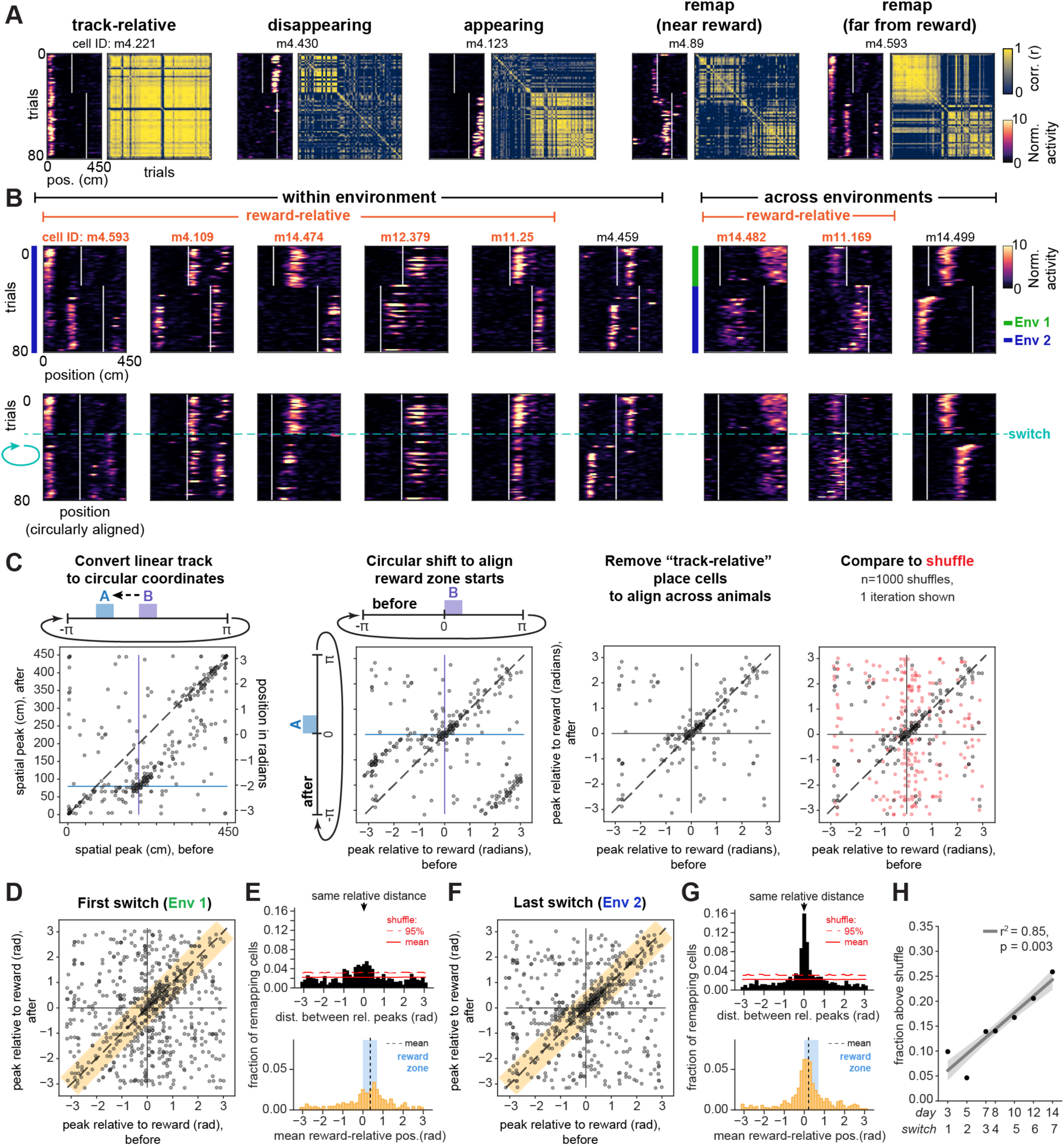
A subpopulation of CA1 cells remaps relative to reward. **A)** Five place cells from an example mouse (m4) with fields that are either track-relative, disappearing, appearing, following the reward location closely (“remap near reward”, with spatial firing peaks ≤ 50 cm from both reward zone starts), or “remap far from reward” (peaks >50 cm from at least one reward zone start; see Methods). Data are from an example switch day in Env 2 (day 14). Left of each cell: Spatial activity per trial shown as smoothed deconvolved calcium events normalized to the mean activity for each cell within session. White lines indicate beginnings of reward zones. Right of each cell: Trial-by-trial correlation matrix for that cell. **B)** Example place cells that remap to the same relative distance from reward (orange cell ID labels), compared to cells that do not (black cell ID labels). Top row: Mean-normalized spatial activity plotted on the original track axis. Bottom row: Post-switch trials (after the horizontal blue dashed line) are circularly shifted to align the reward locations, to illustrate alignment of fields at a similar relative distance from reward both within (left, examples from day 14) and across environments (right, day 8). **C)** Illustration of method to quantify remapping relative to reward. Left: A scatter of peak activity positions before vs. after the reward switch in linear track coordinates reveals track-relative place cells along the diagonal (dashed line; shown for example animal m12 on day 14, switching from reward zone “B” before to “A” after). An off- diagonal band at the intersection of the starts of reward zones suggests cells remapped relative to both rewards. Middle left: We center this reward-relative band by converting the linear coordinates to periodic and rotating the data to align the starts of reward zones at 0. Middle right: Track-relative place cells are removed to isolate remapping cells. Right: The data are then compared to a random remapping shuffle (one shuffle iteration shown in red). Together, these shuffles (n=1000) produce the distribution marked in red lines in (E) and (G). All points are jittered slightly for visualization (see Methods) and are translucent; darker shades indicate overlapping points. **D)** Peak activity positions relative to reward for all animals on the first switch (day 3), for remapping cells with significant spatial information both before and after the switch (n=677 cells, 7 mice). Orange shaded region indicates a range of ≤50 cm (∼0.698 radians) between relative peaks, corresponding to points included in the histogram in (E, bottom). **E)** The fraction of cells remapping to a consistent reward-relative position (≤ 50 cm between relative peaks) on the first switch day is higher than expected by chance. Top: histogram of the circular distance between relative peaks after minus before the switch (i.e. orthogonal to the unity line in D), compared to the mean and upper 95% of the shuffle distribution (solid and dotted red lines, respectively). Bottom: Distribution of mean firing position relative to reward for cells with ≤ 50 cm distance between relative peaks, corresponding to orange shaded area along the unity line in (D) (n=261 cells, 7 mice, non-zero mean = 0.358 radians, 95% confidence interval [lower, upper] = [0.196, 0.519], circular mean test). Black dotted line marks the mean, blue shading marks the extent of the reward zone. **F)** Same as (D) but for the last switch (day 14) (n=707 cells, 7 mice). **G)** Same as (E) but for the last switch (day 14). Bottom: n=398 cells with ≤ 50 cm distance between relative peaks (non-zero mean = 0.206 radians, 95% confidence interval [lower, upper] = [0.103, 0.309], circular mean test). **H)** Fraction of cells above chance that remap relative to reward grows linearly with task experience (number of switches). Each dot shown is the combined fraction across n=7 mice; mean ± confidence interval of linear regression is shown in gray. See also Figure S2.

### An above-chance fraction of reward-relative remapping cells

We next investigated whether a subpopulation of cells encoded experience anchored to reward. To test this hypothesis, we attempted to identify neurons with fields far from the start of the reward zone (>50 cm) that shifted their place fields to match the shift in reward location. To visualize this form of remapping, we circularly shifted the spatial activity of cells on trials following the switch to align the reward locations (**Fig. 2B**). This approach revealed that a subset of cells remapped to a similar relative distance from the reward, both within and across environments (“reward-relative” cells; see below and Methods for full criteria) (**Fig. 2B**). This field alignment relative to reward was observed even for fields that, from the perspective of the linear track position, remapped from the latter half of the track to the beginning, or vice versa (e.g. **Fig. 2B** cells m4.109, m14.482). Such alignment occurred despite the variable length of the teleport zone, suggesting that these cells do not simply track distance run from the last reward. Consistent with this interpretation, the distance run in the teleport zone did not predict the cells’ spatial firing variability on the following lap in the vast majority (∼92%) of reward-relative cells (**Fig. S2E-F**). Instead, reward-relative cells seemed to maintain their preferred firing position with respect to reward according to where the animal was in the progression of the overall task, which spans across the teleport and the trial start but is informed by these boundaries. Altogether, these observations suggest that a reward-relative subpopulation maps a periodic axis of experience from reward to reward, consistent with the cyclical manner in which the animal repeats the task structure. At the same time, other cells seemed to remap randomly, as circular shifting did not align their fields (“non-reward-relative remapping”, 12.3% ± 3.1% of all place cells; **Fig. 2B**, black cell ID labels).

To assess whether reward-relative remapping occurred at greater than chance levels, we plotted the peak spatial firing of all cells that maintained significant spatial information before versus after the switch. This revealed the track-relative place cells along the diagonal, as they maintained their peak firing positions pre- to post-switch. In addition, this analysis revealed an off-diagonal band of cells at the intersection of the reward locations, which extended linearly away from the cluster of reward cells that has been previously described^54^ (**Fig. 2C**, far left panel; **Fig. S2B**, middle). As noted above, we reasoned that the animals may experience the task as periodic instead of a sequence of discrete linear episodes. We therefore transformed the linear track coordinates to the phase of a periodic variable (i.e. 0 to 450 cm becomes -π to π radians), allowing us to shift the data to align the reward zones at zero and isolate the band of cells with peaks at this intersection (**Fig. 2C**, middle-left panel). Cells which fall along the unity line in this analysis thus putatively remap to the same relative distance from reward. We then excluded the track-relative cells to focus on the remapping cells across animals (**Fig. 2C**, middle-right panel). We measured the circular distance of each cell’s peak firing position relative to the reward zone pre- or post-switch (**Fig. 2D**) and calculated the difference between each of these relative peaks. A difference between relative peak positions close to zero indicates reward-relative remapping. We compared the distribution of differences to a “random-remapping” shuffle (**Fig. 2C**, far right panel) and found that the fraction of place cells that exhibited reward-relative remapping across animals exceeded chance on the first switch day (**Fig. 2E**, top). Further, the fraction of cells that exceeded the chance level increased by the last switch day (**Fig. 2F-G, Fig. S2G**) and significantly increased linearly across task experience with each switch (**Fig. 2H**). A linear mixed effects model accounting for mouse identity revealed a similar, significant increase in reward-relative remapping over experience (**Fig. S2H**). Across days and animals, the fraction of cells exceeding chance spanned a mean range of ∼0.65 radians around the unity line, or nearly 50 cm, suggesting there is some variability in the precision of reward-relative remapping (**Fig. 2E, G, Fig. S2G**). This distance thus provided a candidate region to quantitatively identify reward-relative remapping cells. We next analyzed the distribution of reward-relative peak firing positions within this candidate zone (i.e. position along the unity line in **Fig. 2D**) and observed that the mean of the distribution was significantly greater than zero (**Fig. 2E**, bottom). This indicates that more reward-relative place fields are in locations following reward delivery (as indicated by maximal licking, **Fig. 1F-G, Fig. S1A-C**) than in locations preceding the reward. This post-reward shift of the distribution of cells was consistent across days (**Fig. 2G**, bottom).

Next, to investigate the degree to which cells active in close proximity to the reward contribute to the above-chance levels of reward-relative remapping, we excluded cells with peaks within ± 50 cm of the start of both reward zones. Note that this excludes both the reward and anticipatory zones. With this exclusion, we still found above-chance reward-relative remapping (**Fig. S2I-L**). In addition, the mean of the remaining reward-relative firing positions continued to follow the reward zone (**Fig. S2J, L**, bottom). However, there was not a significant increase in remapping at these distances across days (**Fig. S2M**), suggesting there is more growth in the population of reward-relative cells at closer proximities to reward. Nevertheless, this analysis confirmed that reward-relative remapping is not limited to neurons with close proximity to reward.

Finally, we implemented two main criteria to identify a robust subpopulation of reward-relative cells for further analysis. First, we identified candidate place cells with reward-relative spatial peaks that were within 50 cm of each other before versus after a reward switch (i.e. within the orange candidate zone highlighted in **Fig. 2D, F**). This criterion included cells with peaks both very close to and potentially very far from the reward. Second, to reduce the influence of noise in spatial peak detection and take into account the shape of the firing fields, we performed a cross-correlation of the cells’ activity aligned to the reward zone pre- vs. post- switch. We required the offset at the maximum of this cross-correlation to be ≤50 cm from zero lag and exceed chance levels relative to a shuffle for each cell (**Fig. S2N-P**). This approach provides a metric for how stable each cell’s mean firing position is relative to reward. We confirmed that these criteria identified a subpopulation of cells whose fields were maximally correlated in periodic coordinates relative to reward, in contrast to the track-relative cells which were maximally correlated relative to the original linear track coordinates (**Fig. S2O-P**). From here on, we refer to cells that passed both of these criteria as “reward-relative” (RR) (15.4% ± 5.3% of place cells averaged over all switch days; 12.4% ± 3.3% on the first switch, 18.9% ± 6.5% on the last switch).

### Reward-relative sequences are preserved when the reward moves

Next, to ask whether the reward-relative cells constructed sequences of activity anchored to reward, we sorted the reward-relative cells within each animal by their peak firing position on trials before the reward switch, using split-halves cross-validation (see Methods). We applied this sort to the post-switch trials and found that the reward-relative subpopulation indeed constructed sequences that strongly preserved their firing order relative to reward across the switch within an environment (**Fig. 3A-B, Fig. S3A**). Sequence preservation was quantified as a circular-circular correlation coefficient between the peak firing positions in the pre- and post-switch sequences, which was much higher than the shuffle of cell identities for nearly all animals and days (48 out of 49 sessions positively correlated, with rho >95% of the shuffle, two-sided permutation test) (**Fig. 3C**). These sequences spanned the task structure from reward to reward, such that cells firing at the furthest distances from reward would occasionally “wrap around” from the beginning to the end of the linear track or vice versa (e.g. **Fig. 3A**, mouse 4 and **Fig. S3A**). When we introduced the novel environment on day 8, despite apparent global remapping in the total population of place cells (**Fig. S3B**), reward-relative sequences were clearly preserved (**Fig. S3C**, top). The track- relative place cells likewise constructed robust sequences (48 out of 49 sessions positively correlated, with rho >95% of the shuffle, two-sided permutation test) both within (**Fig. 3D-F**) and even across environments (**Fig. S3C**, middle) in a minority of place cells (20.2% ± 8.8% of place cells within vs. 8.4% ± 3.6% across, mean ± s.d. across 7 mice). The sequence stability of both the reward-relative and track-relative subpopulations contrasted starkly with the absence of sequence preservation for most of the non-reward-relative remapping cells (40 out of 49 sessions did not exceed the shuffle; **Fig. 3G-I; Fig. S3C**, bottom) and for the disappearing and appearing cells (**Fig. S3D-E**), as expected.

**Figure 3.**
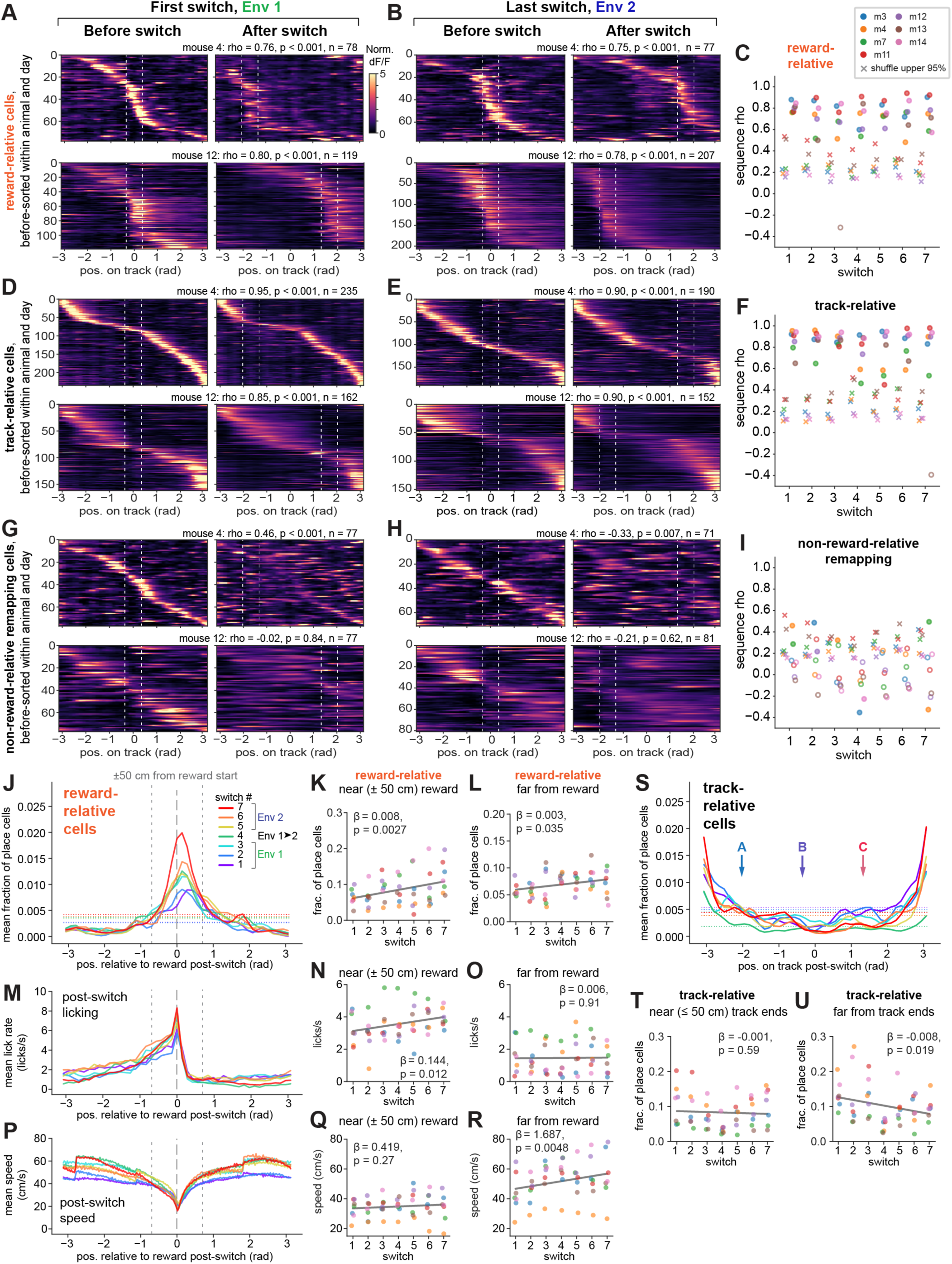
Preserved sequences relative to reward and space. **A)** Preserved firing order of reward-relative sequences across the 1st switch (day 3) (2 example mice, top and bottom rows; m4 is the same mouse shown in Fig. 2A, m12 had the maximum number of cells). Reward-relative remapping cells within each animal are sorted by their cross-validated peak activity before the switch, and this sort is applied to activity after the switch. Data are plotted as the spatially-binned, unsmoothed dF/F normalized to each cell’s mean within session, from which peak firing positions are identified in linear track coordinates that have been converted to radians (see Fig. 2C). White dotted lines indicate starts and ends of each reward zone. Each sequence’s circular-circular correlation coefficient (rho), p-value relative to shuffle, and number of cells is shown above, where p < 0.001 indicates that the observed rho exceeded all 1000 shuffles. Color scale is applied in (B), (D-E), (G-H). **B)** Reward-relative sequences on the last switch (day 14), for the same mice shown in (A). **C)** Circular-circular correlation coefficients for reward-relative sequences before vs. after the reward switch, for each mouse (n=7), with the upper 95^th^ percentile of 1000 shuffles of cell IDs marked as “x”. Closed circles indicate coefficients with p < 0.05 compared to shuffle using a two-sided permutation test; open circles indicate p ≥ 0.05. **D)** Sequences of track-relative place cells on the 1st switch day for the same mice as in (A-B). **E)** Sequences of track-relative place cells on the last switch day for the same mice as in (A-B,D). **F)** Circular-circular correlation coefficients for track-relative place cell sequences (n=7 mice), colored as in (C). **G-H)** Sequences of non-reward-relative remapping cells with significant spatial information before and after the switch, for the same mice as above and on the 1st (G) and last (H) switch day. Note that a cross-validated sort of the “before” trials produces a sequence, but in general this sequence is not robustly maintained across the switch. **I)** Circular-circular correlation coefficients for non-reward-relative remapping cells (n=7 mice), colored as in (C). **J)** Distribution of peak firing positions for the reward-relative sequences after the switch on each switch day, shown as a fraction of total place cells within each animal, averaged across animals (n=7 mice). Standard errors are omitted for visualization clarity. Horizontal dotted lines indicate the expected uniform distribution for each day (see Methods); vertical gray dashed lines indicate the reward zone start and surrounding ±50 cm. **K-L**) Quantification of changes in reward-relative sequence shape across animals and days, using linear mixed effects models with switch index as a fixed effect and mice as random effects (n=7 mice). β is the coefficient for switch index, shown with corresponding p-value (Wald test). Each point is the fraction of place cells within that mouse (colored as in C) in the reward-relative sequences, specifically within ±50 cm (K) or >50 cm (L) from the start of the reward zone. **M)** Mean lick rate across post-switch trials and across animals (10 cm bins), colored by switch as in (J). **N-O)** Quantification of change in lick rate across days within ±50 cm of the reward zone start (N) or >50 cm (O), using a linear mixed effects model as in (K-L). **P)** Mean running speed across post-switch trials and across animals (2 cm bins), colored by switch as in (J). **Q-R)** Quantification of change in speed across days, as in (K-L) and (N-O). **S)** Distribution of peak firing positions for the track-relative sequences after the switch on each switch day, shown on the track coordinates converted to radians (not reward-relative). Arrows indicate the start position of each reward zone (“A”, “B”, “C”). Horizontal dotted lines indicate the expected uniform distribution for each day. **T-U)** Quantification of changes in track-relative sequence shape across animals and days as in (K-L), here within ≤50 cm of either end of the track (T) or >50 cm from the track ends (U). See also Figure S3.

We next quantified the mean sequence shape for each subpopulation, either relative to reward or to the VR track coordinates. The reward-relative sequences strongly over-represented the reward zone^1,2,54,55,63–66^ and were most concentrated just following the start of the reward zone (**Fig. 3J**). Further, this density near the reward zone grew across task experience (**Fig. 3J-K**). The fraction of place cells at further positions (>50 cm) from the start of the reward zone showed a modest increase over days, but to a much lesser degree than positions near the reward (**Fig. 3L**). In parallel, mice increased their average post-switch lick rate prior to the reward zone (**Fig. 3M-O**) and their running speed away from the reward zone over days (**Fig. 3P-R**). These changes suggest that reward-relative sequence strength increases as behavioral performance becomes more robust. In addition, on any given day, the precision of the animal’s licking correlated with the precision of the reward-relative sequence (i.e. the narrowness of the cell distribution around reward, measured as the circular variance) (**Fig. S3F-I**). In contrast to the reward- relative sequences, the track-relative place cell sequences tended to over-represent the ends of the track, as opposed to the reward zones (**Fig. 3S**). In addition, the track-relative sequences showed a subtle but significant drop in density at positions away from the ends of the track across days, consistent with selective stabilization of place fields near key landmarks^8,55,76–80,86^ (**Fig. 3S-U**). These differences in sequence shape between the reward-relative and track-relative cells suggest that each of these subpopulations anchors to the most salient point in the “space” they encode. For the reward-relative sequences, this is the reward zone, and for the track-relative sequences, this is likely the start and end of the track where the animal exits and enters the teleport zone. By comparison, the disappearing, appearing, and non-reward-relative remapping cells encoded a more uniform representation of the track (**Fig. S3J- K**), with only subtle over-representation of the track ends and an observable decrease across the track for the disappearing cells across days (**Fig. S3J-K**).

### Non-track-relative place cells are increasingly recruited to the reward-relative population over days

The growth in reward-relative sequence density across task experience raised three possibilities: (1) reward-relative cells may become more consistent at remapping to precise reward-relative positions with extended learning, (2) more cells might be recruited from other populations to the reward-relative population, or (3) a combination of both changes occurs. To investigate these possibilities and understand how individual neurons are dynamically reallocated to represent updated reward information within and across environments, we followed the activity of the same neurons across days by matching the aligned regions of interest (ROIs) for each cell. Specifically, we followed single neurons across pairs of switch days and analyzed their spatial firing patterns from one switch to the next (**Fig. 4**). Using linear mixed effects models to account for variance across animals, we found that there was an increase in the fraction of cells that remained reward-relative on consecutive reward switches, indicating that reward-relative cells are more likely to maintain their reward-relative coding across experience (**Fig. 4A-B**). In addition, an increasing proportion of non-track-relative cells (appearing, disappearing, and non-reward-relative remapping) were recruited into the reward-relative population with task experience (**Fig. 4C-D**). Non- place cells and track-relative cells likewise were recruited into the reward-relative population, though this occurred at a constant rate of turnover that did not increase over days (**Fig. 4E-H**). In addition, the track- relative cell population did not recruit more cells from any other population (**Fig. 4I-K**) and did not show an increased likelihood of remaining track-relative across experience (**Fig. 4L**), in contrast to the reward- relative population. These results suggest that the track-relative place cells remain a more independent population from the reward-relative cells, and that cells exhibiting high flexibility (such as appearing/disappearing) are more likely to be recruited into the reward-relative representation with extended experience in this task.

**Figure 4.**
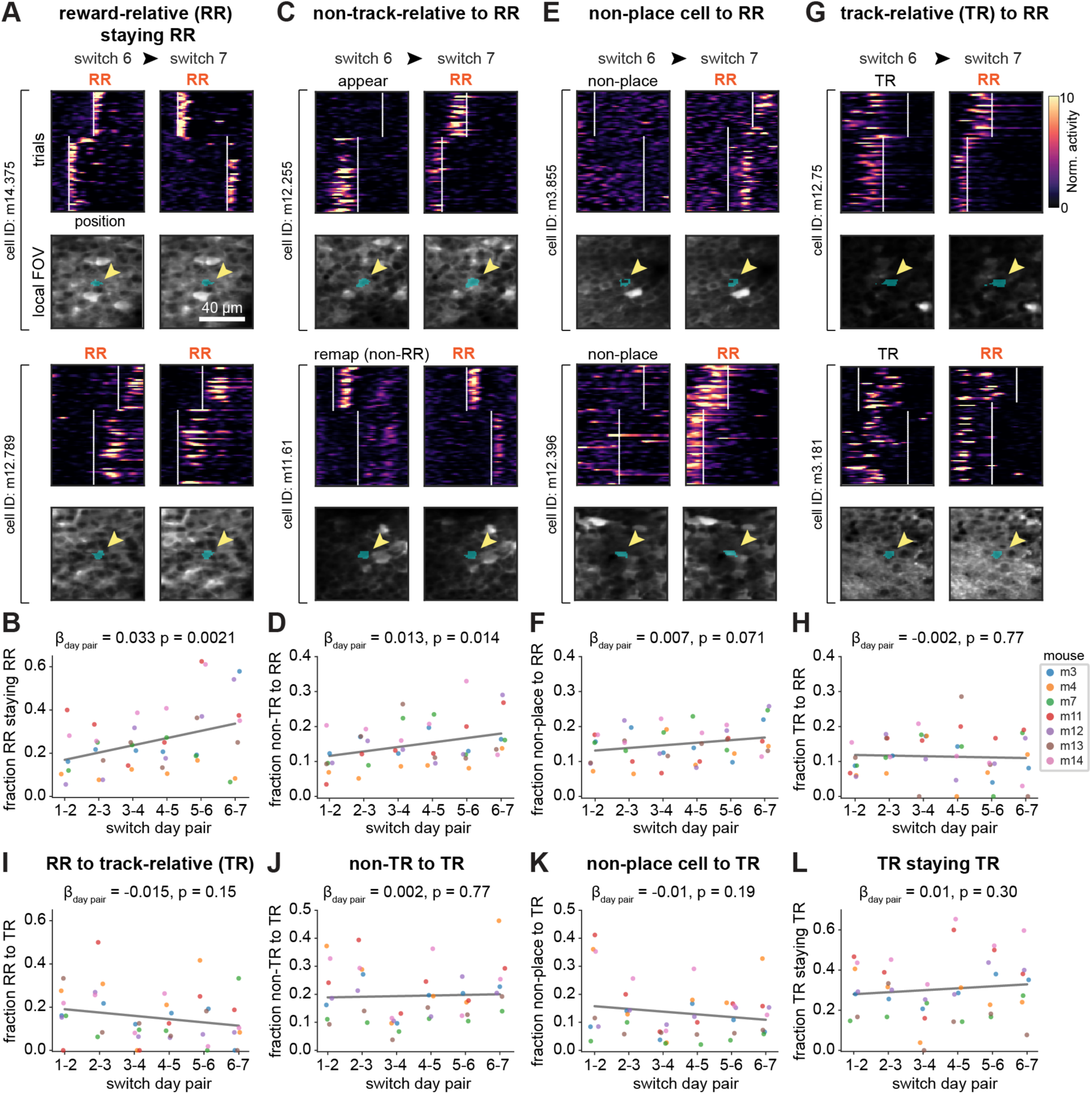
Tracking cells over days reveals increased recruitment into the reward-relative population. **A)** Example cells from 2 separate mice that were tracked from switch 6 to 7 (days 12-14) that were identified as reward-relative (RR) on both switch 6 and switch 7, and which maintained their firing position relative to reward. Top of each cell: spatial firing over trials and linear track position on each day (deconvolved activity normalized to the mean of each cell). White vertical lines indicate the beginnings of the reward zones. Bottom of each cell: Mean local field of view (FOV) around the ROI of the cell on both days (blue shading highlights ROI, indicated by the yellow arrow). **B)** Fraction of tracked RR cells on each switch day that remain RR on subsequent switch days increases over task experience. All β coefficients and p-values in (B, D, F, H, and I-L) are from linear mixed effects models with switch day pair as a fixed effect and mice as random effects (n=7 mice in each plot, key in H). Coefficients correspond to the fractional increase in recruitment per switch day pair, from the model fit (gray line). **C)** Same as (A) but for 2 example cells that were identified as appearing (top) or non-RR remapping (bottom) on switch 6 but were converted to RR on switch 7. **D)** Fraction of “non-track-relative” place cells (combined appearing, disappearing, and non-RR remapping) that convert into RR cells on subsequent switch days increases over task experience. **E)** Same as (A) but for 2 example non-place cells on switch 6 (did not have significant spatial information in either trial set) that were converted to RR place cells on day 7. **F)** Fraction of non-place cells converting to RR cells on subsequent switch days shows a modest increase over task experience. **G)** Same as (A) but for 2 example track-relative (TR) cells on switch 6 that were converted to RR on switch 7. **H)** Fraction of TR cells converting to RR cells does not increase over task experience. **I)** Fraction of RR cells converting to TR cells does not change significantly over task experience. **J)** Fraction of non-TR cells (combined appearing, disappearing, and non-RR remapping) converting to TR does not change significantly. **K)** Fraction of non-place cells converting to TR does not change significantly. **L)** Fraction of TR cells remaining TR does not change significantly. See also Figure S4.

The track-relative population itself showed a steady rate of turnover over days (**Fig. 4L**), consistent with representational drift^81–85^. Indeed, when we analyzed place cell sequences identified on each reference day and followed two days later, we observed drift even in the fixed-condition animals (n=3 mice) that experienced a single environment and reward location for all 14 days (**Fig. S4A, F**). Drift was prevalent as well in the reward-relative and track-relative sequences tracked across consecutive switch days (**Fig. S4B-E, G-J**), at a slightly higher degree than the fixed-condition animals (**Fig. S4F**), consistent with our observation that moving a reward induces more remapping than not moving a reward (**Fig. 2A, Fig. S2A-D**). In a subset of animals and day pairs, cells that remained reward-relative or track-relative tended to maintain their firing order across days. However, sequence preservation across days was generally lower (i.e. closer to the shuffle) (**Fig. S4C, E, H, J**) than within-day sequences (**Fig. 3C, F; Fig. S3A, C**). These results indicate that rather than being dedicated cell classes, reward-relative and track- relative ensembles are both subject to the same plasticity across days of experience as the overall hippocampal population^81–85^. However, the increasing recruitment of cells into the reward-relative population, despite this population having different membership on different days, suggests an increased allocation of resources to the reward-relative representation. We conclude that as animals become more familiar with the task demands—specifically the requirement to estimate a hidden reward location that can be dissociated from spatial stimuli—the hippocampus builds an increasingly dense representation of experience relative to reward, while preserving a fixed representation of the spatial environment in a largely separate population of cells on any given day.

### Encoding of reward proximity versus movement covariates in reward-relative cells

The animals’ approach to and departure from the reward zone in this task is associated with stereotyped running and licking behaviors that comprise an important aspect of the animal’s experience surrounding reward. For instance, these behaviors must be timed with an accurate estimate of distance from reward to inform behavioral adaptations after the reward switch. We therefore took two complementary approaches, detailed below, to disentangle which features of the reward-relative experience are encoded in the reward-relative subpopulation. These approaches further allowed us to control for the potential influence of movement covariates on the neural activity.

First, we tested the hypothesis that reward-relative firing following the reward could be locked to the running speed profile of the animal, as opposed to the location where the animal expected to receive reward. For this, we leveraged the ∼15% omission trials to compare the activity of reward-relative neurons on rewarded versus unrewarded trials. To consider the activity of reward-relative cells under conditions where running speed was generally comparable between rewarded and omission trials, we restricted our analysis to: reward-relative cells with peaks following the start of the reward zone, sessions in which the reward zone began at location “A” (80 cm) or “B” (200 cm), and trials before a reward switch. To then align the running speed profiles across rewarded and omission trials, we fit a time warping model^87^ on the running speed data and applied this time warped transformation to the neural data (**Fig. S5A-D**, see Methods). We then computed a “reward versus omission index” (RO index) to quantify the difference in each cell’s firing rate following rewards versus omissions (**Fig. S5E**).

Reward-relative cells exhibited heterogeneity in their firing rates following rewards versus omissions (**Fig. 5A-C**). A subset of cells fired at the same position relative to reward regardless of the animal’s running speed (**Fig. 5A**, top and middle row of cells, **Fig. 5B**, middle, **Fig. 5C**, top and middle), with firing rates often differing between rewarded and omission trials (e.g. **Fig. 5A**, top, **Fig. 5C**, top and middle). Another subset of cells fired relative to a particular phase of the speed profile (**Fig. 5A**, bottom, **Fig. 5B**, top and bottom, **Fig. 5C**, bottom). However, at the population level, the distribution of RO indices for reward-relative cells was skewed toward reward-preferring (**Fig. 5D**). This reward-preference was also clearly observable at the single-cell and population levels in trials after the reward switch (**Fig. S5F-I**), although running speed is also more variable on these trials (**Fig. S5F**). Finally, combining RO indices across days (**Fig. S5J**) revealed that, while there is heterogeneity among individual cells, there is a significant preference for the reward-relative population to fire more on rewarded rather than omission trials (**Fig. 5E, Fig. S5K**; note results were similar when all sessions were included, **Fig. S5L**).

**Figure 5.**
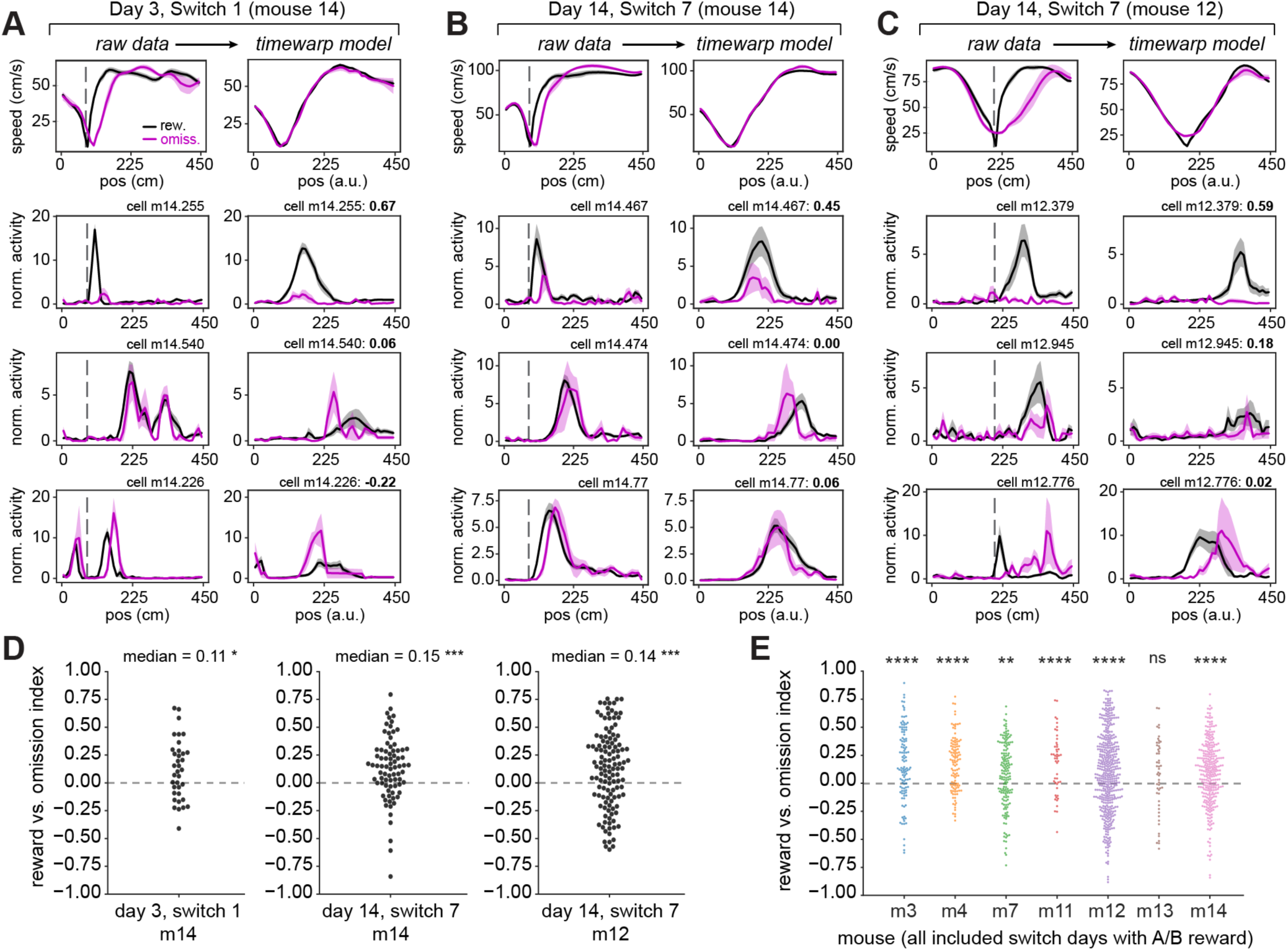
Diversity in reward-relative cell coding on rewarded vs. omission trials, with a population preference for reward. **A)** Raw and time warp model-aligned speed (first row) and neural activity for 3 reward-relative example cells (second to fourth rows) on the first switch day, in mouse 14. Throughout Fig. 5, analysis is performed only on trial sets before the reward switch which have at least 3 omission trials. In (A-C), the left columns show raw (untransformed) data, with the start of the reward zone indicated by the gray dashed line. The right columns show data transformed by the time warp model fit to the trial-by-trial speed in that session (see Methods, Fig. S5). All included cells have peak firing following the start of the reward zone. First row: mean ± s.e.m. speed profiles for rewarded (black) and omission (magenta) trials. Note precise alignment of the transformed speed profiles on the right. Second to fourth rows: mean ± s.e.m. deconvolved activity for rewarded and omission trials, normalized to each cell’s mean within the session, showing an example reward-relative cell in each row. The difference in model-transformed activity between rewarded and omission trials is captured as a “reward vs. omission” index (bold text, top right of each panel; see Methods). The top row highlights neurons with much higher activity on rewarded trials. A positive index indicates higher activity on rewarded trials, a negative index indicates higher activity on omission trials. **B)** Raw and time warp model-aligned speed and neural activity for 3 reward-relative example cells on the last switch day, in mouse 14. Note cell m14.474 is also shown in Fig. 2B. **C)** Raw and time warp model-aligned speed and neural activity for 3 reward-relative example cells on the last switch day, in mouse 12. Note that speed is still aligned well with the reward zone at position “B” (at 220 cm) for m12 vs. position “A” (at 80 cm) for m14. Note cell m12.379 is also shown in Fig. 2B. **D)** Reward vs. omission index for the population of reward-relative cells that have spatial firing peaks following the start of the reward zone, in the session shown in (A) (left), (B) (middle), and (C) (right). Each dot is a cell. *p<0.05, ***p<0.001, one-sample Wilcoxon signed-rank test against a 0 median. Switch 1, m14: W = 183, p = 0.018, n = 36 cells; Switch 7, m14: W = 721, p = 2.69e-5, n = 79 cells; Switch 7, m12: W = 2358, p = 3.70e-4, n = 122 cells. **E)** Reward vs. omission index across all switch days where the reward zone was at “A” or “B”, for the population of reward-relative cells that have spatial firing peaks following the start of the reward zone within each mouse (n = 7 mice). Each dot is a cell, colors correspond to individual mice. Significance level is set at p<0.007 with Bonferroni correction. **p<0.007, ****p<0.0001. m3: median = 0.14, W = 1.50e3, p = 4.66e-7, n = 3 days, 114 cells; m4: median = 0.20, W = 1.03e3, p = 5.93e-10, n = 3 days, 112 cells; m7: median = 0.09, W = 4.44e3, p = 2.08e-3, n = 4 days, 157 cells; m11: median = 0.23, W = 194, p = 5.32e-5, n = 4 days, 48 cells; m12: median = 0.09, W = 3.41e4, p = 4.40e-8, n = 5 days, 441 cells; m13: median = 0.09, W = 523, p = 0.13, n = 3 days, 52 cells; m14: median = 0.11, W = 1.39e4, p = 3.74e-8, n = 4 days, 296 cells. See also Figure S5.

Second, we implemented a generalized linear model (GLM)^88^ to dissociate the contribution of task versus movement variables to the moment-to-moment deconvolved calcium activity of each cell. For task variables, we included linear track position, reward-relative position, and whether the animal was rewarded on each trial, convolved with the linear track position. For movement variables, we included running speed, acceleration, and licking (**Fig. S6A**, see Methods for details). We trained the GLM using 5-fold cross-validation and tested its prediction on the activity of each neuron in held out trials (**Fig. 6A- B, Fig. S6B-C**). Only well-fit neurons (fraction deviance explained > 0.15)^88^ were included in the analysis (35% of place cells across 7 mice and 7 switch days; **Fig. S6B-C**). This included 61% (2377/3901) of track-relative cells, 37% (1157/3104) of reward-relative cells (**Fig. 6A-B**), and 43% (1091/2518) of non- RR remapping cells.

**Figure 6.**
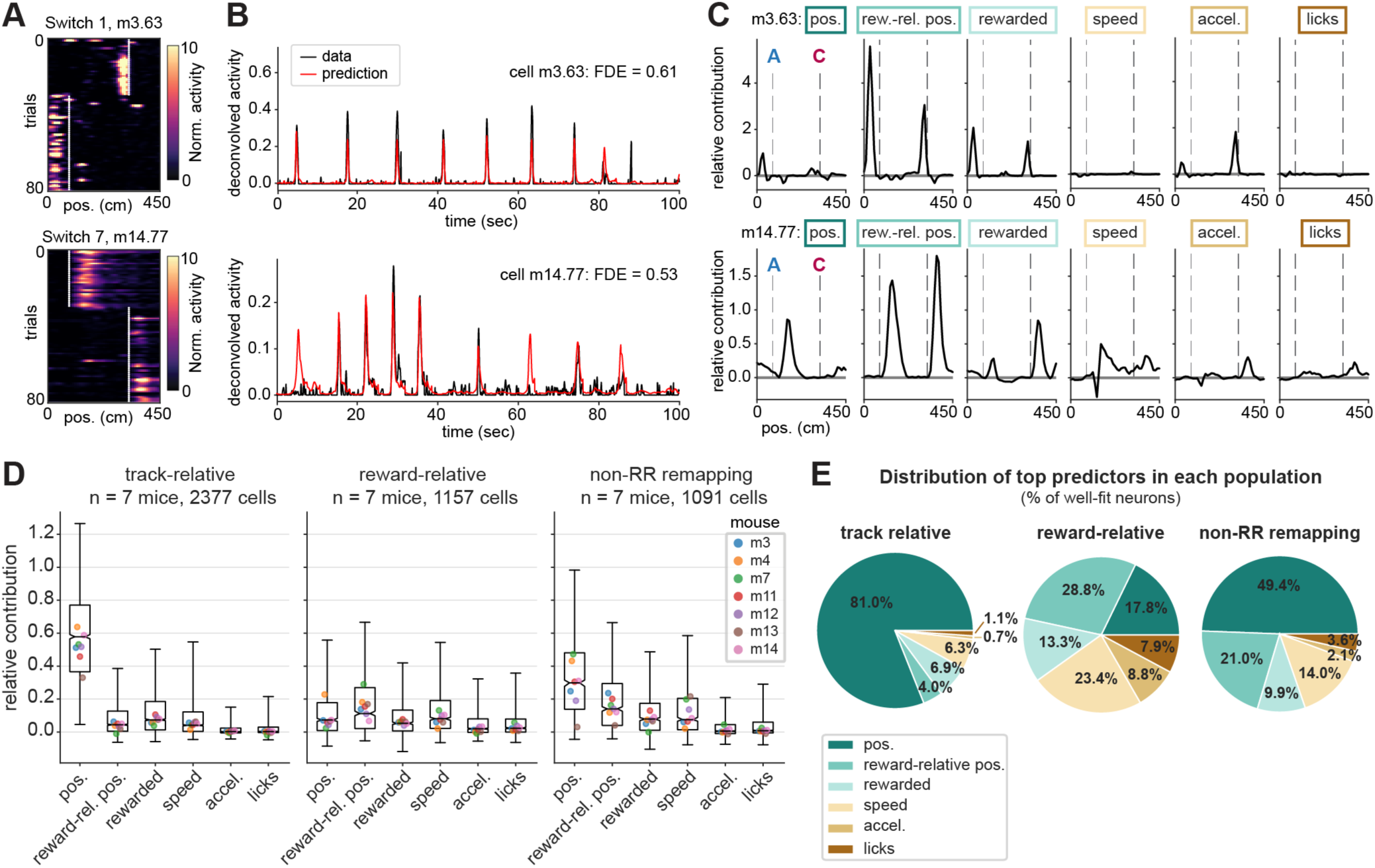
A generalized linear model reveals reward-relative position as a leading predictor of neural activity in the reward-relative population. **A)** Example spatial firing maps of two reward-relative cells that were fit well (fraction deviance explained > 0.5) by the GLM, on the first switch day (top) and last switch day (bottom). White lines mark the starts of reward zones. Note that cell m14.77 (bottom) is the same cell as shown in Fig. 5B, bottom row. **B)** Deconvolved calcium activity (black) and GLM prediction (red) of held-out test data for the two example cells shown in (A). Fraction Poisson deviance explained (FDE) of the test data is reported as a measure of model performance. **C)** Relative contribution (“fraction explained deviance”, see Methods), or encoding strength, for each variable included in the GLM binned by track position, for the two example cells shown in (A). Task variables are “pos.”: linear track position; “rew.-rel. pos.”: reward-relative position; “rewarded”: whether the animal was rewarded as a function of position (i.e. a binary that stays high from the reward delivery time to the end of the trial). Movement variables are speed, acceleration, and licking (see Fig. S6A for implementation). Relative contribution is calculated from an ablation procedure where each variable is individually removed from the full model (see Methods). Gray dashed lines mark the starts of the two reward zones in each session, also indicated in (A). Boxes correspond to color coding in (E). Note that reward-relative position provides the maximum relative contribution for both example cells, including m14.77 even though it fired equivalently on rewarded and omission trials (see Fig. 5B). **D)** Distributions of relative contribution of each variable in the track-relative, reward-relative, and non-reward- relative remapping subpopulations across animals. Boxes indicate the interquartile range, whiskers extend from the 2.5th to 97.5th percentile, horizontal line indicates the median, and notches indicate the confidence interval around the median computed from 10,000 bootstrap iterations. Colored dots mark the medians of each individual mouse. **E)** Distributions of top predictor variables (maximum relative contribution) for individual cells within each subpopulation, shown as % of cells in the subpopulation; same n as in (D). See also Figure S6.

After testing the full model, we performed an ablation procedure to measure the relative contribution of each predictor variable (see Methods). This ablation procedure revealed clear peaks in the contribution of reward-relative position to the activity of many reward-relative cells as a function of position along the track, recapitulating their firing fields (**Fig. 6C**). We then quantified the relative contribution of each variable within the track-relative, reward-relative, and non-RR remapping subpopulations that were well-fit by the full model (**Fig. 6D**). We found that linear track position was the strongest predictor for the track-relative population (**Fig. 6D**) and the top predictor for 81% of included track-relative cells (**Fig. 6E**). This result provided confirmation of our classification of these cells as stable place cells that remain locked to the spatial environment within a session. The non-RR remapping cells likewise were best predicted by linear track position followed by reward-relative position (**Fig. 6E**), consistent with their recruitment to the reward-relative population over experience (**Fig. 4**). In contrast, for the reward-relative population, reward-relative position provided the highest relative contribution (**Fig. 6D**) and the top predictor for 28.8% of reward-relative cells (**Fig. 6E**). Reward-relative position was followed by speed (top predictor for 23.4% of reward-relative cells), linear track position (17.8%), and the receipt of reward (13.3%), with minimal contributions of acceleration and licking. Within the reward-relative cells that exhibited reward-relative position or speed as their top predictors, we observed a mixed contribution of the other variables as well (**Fig. S6D-E**). These results suggest that the reward-relative population supplies a heterogeneous code for multiple aspects of the reward-related experience. While movement covariates are challenging to fully disentangle from the animal’s progression through the task, the GLM, combined with our analyses of rewarded versus omission trials, provides evidence that reward and reward-relative position are strong predictors of activity in the reward-relative cell population.

## DISCUSSION

Here, we reveal that the hippocampus encodes an animal’s physical location and its position relative to rewards through parallel sequences of neural activity. We employed 2-photon calcium imaging to observe large groups of CA1 neurons in the hippocampus as mice navigated virtual linear tracks with changing hidden reward locations. This paradigm allowed us to identify reward-relative sequences that are dissociable from the spatial visual stimuli, as they instead remain anchored to the movable reward location. This suggests that the brain employs a parallel coding system to simultaneously track an animal’s physical and reward-relative positions, pointing to a mechanism for how the brain could monitor an animal’s experience relative to multiple frames of reference^45,59,60,67–73,89^. Further, we found that the reward-relative sequences, but not the “track-relative” sequences locked to the spatial environment, show a notable increase in the number of neurons involved as the animal gains more experience with the task. These results suggest that the hippocampus builds a strengthening representation of generalized events surrounding reward as animals become more familiar with the task structure. Moreover, this work raises the possibility that the hippocampal code for space, provided by a subpopulation of place cells, can update the progression of reward-relative sequences when the reward location changes. In this way, hippocampal ensembles encoding different aspects of experience may interact to support accurate memory while amplifying behaviorally salient features such as reward.

We validated prior work that found a subpopulation of hippocampal neurons precisely encodes reward locations^54^. Here, we found that this subset of reward-relative neurons, which fire very close to rewards, comprised a central component of a more extensive reward-relative sequence. Our finding of reward-relative coding at farther distances from reward is reminiscent of hippocampal cells that encode distances and directions to goals in freely flying bats^90^. At the same time, we extended these previous findings in two key ways: (1) we found that the reward-relative code spans the entire environment, involving cells firing at distances up to hundreds of centimeters away (i.e. as far as from one reward to the next), and (2) additional neurons are recruited into the reward-relative sequences over time, demonstrating a dynamic and evolving coding strategy. Specifically, the cells firing closest to reward were most likely to increase in number across days (**Fig. 2D-H, Fig. S2I-M, Fig. 3J-L**), suggesting that they may be the most detectable in other paradigms. Importantly, we also observed a noticeable turnover in the neurons that form the reward-relative population from day to day, indicating a high degree of reorganization in terms of which neurons participate. Despite this daily fluctuation in individual neurons, the overall sequential structure appears to be maintained within the population.

This latter finding of day to day fluctuation in the reward-relative population is consistent with recent studies which have highlighted a phenomenon known as representational drift^81–85^. In the hippocampus^81–85^ and in other cortical areas^91–93^, this drift is characterized by changing spatial tuning and stability of single neurons as a function of extended experience^82,83^. Despite these single neuron changes, the population often continues to accurately code for the spatial environment^81^ or task^91^. This flexibility is thought to increase the storage capacity of neuronal networks^94–97^. Related changes over time have also been proposed to support the decoding of temporal separation between unique events^50,92,94,98^, which could be particularly important for separating memories of events surrounding different rewarding experiences. We observed population drift in both the reward-relative and track-relative sequence membership. This frequent reorganization suggests that these are not dedicated subpopulations; neurons in the reward-relative population occasionally switched coding modes—though this flexibility may vary by anatomical location within CA1^54,99,100^. At the same time, we found that more resources (i.e. neurons) are allocated to the reward-relative representation at the population level with extended experience. This finding indicates an increasingly robust network code that can be reliably anchored to reward within a day despite day-to-day drift. This stability amidst flux offers insight into the brain’s ability to balance dynamic coding with the preservation of population-level representations^91,92,97^.

In the hippocampus, spatial firing can also be highly flexible at short timescales even within a fixed environment. This flexibility is evident in the variability of place cell firing even within a field^57,85^ and especially when an animal takes different routes^101–106^ or targets a different goal^89,90,107–109^ through the same physical position. Our work is consistent with such trajectory specific coding and raises the possibility that the reward-relative population may split even further by specific destinations or reward sites when there are multiple possible reward locations. Indeed, in cases where multiple rewards are present at once, hippocampal activity may multiplex spatial and reward-relative information to dissociate different goal locations^64,89,110^. This discrimination is enhanced over learning^110^, consistent with the manner in which hippocampal maps are likewise dependent on learned experience^4,6,7,12,52,53,111^.

Furthermore, place cells undergo significant remapping in response to changes in the animal’s motivational state, such as during task engagement^61,62,70^ or disengagement^112^, or upon removal of a reward^113^. Importantly, in our task the animal must remain engaged when the reward zone updates in order to accurately recall and seek out the new reward location. Consistent with this enhanced attention^58,59,61,62^, we see predictable structure in how the neurons remap surrounding reward (i.e. they maintain their firing order relative to reward), in contrast to the more random reorganization observed with task disengagement^112^ or the absence of reward^113^. However, an important consideration for all of these short timescale firing differences, and for representational drift, is the contribution of behavioral variability to neural activity changes^114,115^. For example, small changes in path shape^116^ and the movement dynamics involved in path integration^117–121^ may influence aspects of hippocampal activity. Our results from a generalized linear model indicate that such movement dynamics influence but do not dominate the activity in the reward-relative population, as reward-relative position was a leading predictor of activity. Further work will be required to disentangle which aspects of reward-seeking behavior are encoded in the hippocampus.

The origin of the input that informs reward-relative neurons remains an open question. One possibility is that subpopulations of neurons in CA1 receive stronger neuromodulatory inputs than others^1^, increasing their plasticity in response to reward-related information^122,123^. CA1 receives reward-related dopaminergic input from both the locus coeruleus (LC)^124,125^ and the ventral tegmental area (VTA)^113,126,127^ that could potentially drive plasticity to recruit neurons into reward-relative sequences. Both of these inputs have recently been shown to restructure hippocampal population activity, with both LC and VTA input contributing to the hippocampal reward over-representation^113,125,126^, and VTA input additionally stabilizing place fields^113^ in a manner consistent with reward expectation^113,128^. Alternatively, it is possible that upstream cortical regions, such as the prefrontal cortex^129^, contribute to shaping these reward-relative sequences. Supporting this idea, the prefrontal cortex has been observed to exhibit comparable neural activity sequences that generalize across multiple reward-directed trajectories^130–134^ to encode the animal’s progression towards a goal^135,136^. Another potential source of input is the medial entorhinal cortex, which exhibits changes in both firing rate and the location where cells are active in response to hidden reward locations^137,138^. In particular, layer III of the medial entorhinal cortex is thought to contribute significantly to the over-representation of rewards by the hippocampus^63^. Additionally, the lateral entorhinal cortex, which also projects to the hippocampus, exhibits a strong over-representation of positions and cues just preceding and following rewards^120,139–141^ and provides critical input for learning about updated reward locations^139,140^.

These insights point to a complex network of brain regions potentially interacting to inform reward-relative sequences in the hippocampus. In turn, the hippocampal-entorhinal network may send information about distance from expected rewards back to the prefrontal cortex and other regions where similar activity has been observed^1^, such as the nucleus accumbens^142–147^. Such goal-related hippocampal outputs to prefrontal cortex^148^ and nucleus accumbens^149^ have recently been reported, specifically in the deep sublayer of CA1^148^, where goal-related remapping is generally more prevalent^148,150^. Over learning, reward-relative sequences generated between the entorhinal cortices and hippocampus may become linked with codes for goal-related action sequences in the prefrontal cortex and striatum, helping the brain to generalize learned information surrounding reward to similar experiences^1^. Understanding these interactions in future work will provide new insight regarding how the brain integrates spatial and reward information, which is crucial for navigation and decision-making processes.

The significance of reward-relative sequences in the brain may lie in their connection to hidden state estimation^7,43,151–155^, a concept closely related to generalization^28,156^. Hidden state estimation refers to the process by which an animal or agent infers an unobservable state, such as the probability of receiving a reward under a certain set of conditions, in order to choose an optimal behavioral strategy^151^. For example, in our study involving mice, reward-relative sequences could be instrumental in helping the animals discern different reward states, such as whether they are in a state of the task associated with reward A, B, or C. By building a code during learning that allows the hippocampus to accurately infer such states, this inference process can generalize to similar future scenarios in which the animal must make predictions about which states lead to reward^151,152^. Interestingly, however, the density of the reward-relative sequences we observed was highest following the beginning of the zone where animals were most likely to get reward, rather than preceding it. This finding contrasts with computational simulations of predictive codes in the hippocampus that are skewed preceding goals^157^. Instead, the shape of the reward-relative sequences may be more consistent with anchoring to the animal’s estimate of the reward zone, with decreasing density around the zone as a function of distance which mirrors the organization seen around other salient landmarks^86,158^.

Our work is consistent with previous proposals that suggest a primary function of the hippocampus is to generate sequential activity at a baseline state^17,27,29,47,48^. These sequences could then act as a substrate that can become anchored to particular salient reference points^16,17,27,47^, allowing maximal flexibility to learn new information as it becomes relevant and layer it onto existing representations^159–161^. This role for hippocampal sequences extends to computational studies^28,159,162^ and human research^163–167^, where such coding schemes have been linked to the organization of knowledge and the chronological order of events in episodic memory^27,49,163–166,168–170^. The ability of the hippocampus to parallelize coding schemes for different features of experience may help both build complete memories and generalize knowledge about specific features without interfering with the original memory. As observed in our study, the hippocampus may then amplify the representations of some of these features over others according to task demands. A similar effect has been observed in humans, in that memory recall is particularly strong for information that either precedes or follows a reward^171,172^. This suggests that rewards can significantly enhance the strength and precision of memory storage and retrieval, especially for events closely associated with these rewards. Thus, reward-relative sequences may play a crucial role in how we form and recall our experiences, with implications stretching beyond spatial navigation.

## Author contributions

MS, MHP, and LMG conceptualized the study. MS collected the data and designed and performed analyses. MHP developed data preprocessing and analysis infrastructure, built the microscope and virtual reality rig, and collected data for pilot studies. LMG supervised the project. MS, MHP, and LMG wrote the manuscript.

## Acknowledgements

We thank Charlotte Herber, John Wen, and Jon Rueckemann for comments on the manuscript; Can Dong for assistance with animal training; Adriana Diaz for histology assistance and lab infrastructure support; Alex Gonzalez, Tucker Fisher, and Eric Denovellis for guidance on model implementation and evaluation; and Shih-Yi Tseng for feedback on the GLM. We thank members of the Giocomo laboratory and the Simons Collaboration for the Global Brain Remapping postdoc group for extensive feedback and discussions, in particular P. Dylan Rich, E. Mika Diamanti, and Eric Denovellis for suggesting edits to draft figures. We thank William Newsome and Scott Linderman for frequent feedback on this work, Charline Tessereau and Feng Xuan for insightful discussions, Simón Carrillo-Segura and André Fenton for generously sharing code and valuable insights, and Jeff Gauthier for input on calcium signal processing and discussions about reward-related remapping. This work was supported by a Helen Hay Whitney Foundation fellowship (MS), National Institute of Health Grants 1R01MH126904-01A1 (LMG), R01MH130452 (LMG), BRAIN Initiative U19NS118284 (LMG), P50 DA042012 (LMG), The Vallee Foundation (LMG), The James S. McDonnell Foundation (LMG), The Simons Foundation 542987SPI (LMG), and the Champalimaud Vision Award to William Newsome, who we thank for sharing his support.

## Declarations of Interests

The authors declare no competing interests.

## Data and Code Availability

Upon publication, data will be made available on a publicly accessible data sharing site (e.g. figshare, DANDI, Mendeley). Custom scripts for analyzing the data will be available on GitHub.

## METHODS

### Subjects

All procedures were approved by the Institutional Animal Care and Use Committee at Stanford University School of Medicine. Male and female (n = 5 male, 5 female) C57BL/6J mice were acquired from Jackson Laboratory. Mice were housed in a transparent cage (Innovive) in groups of five same-sex littermates prior to surgery, with access to an in-cage running wheel for at least 4 weeks. After surgical implantation, mice were housed in groups of 1-3 same-sex littermates, with all mice per cage implanted. All mice were kept on a 12-h light–dark schedule, with experiments conducted during the light phase. Mice were between ∼2.5 to 4.5 months at the time of surgery (weighing 18–31 g). Before surgery, animals had *ad libitum* access to food and water, and *ad libitum* access to food throughout the experiment. Mice were excluded from the study if they failed to perform the behavioral pre-training task described below.

### Surgery: calcium indicator virus injections and imaging window implants

We adapted previously established procedures for 2-photon imaging of CA1 pyramidal cells^7,173^. The imaging cannula was constructed using a ∼1.3-mm long stainless-steel cannula (3 mm outer diameter, McMaster) affixed to a circular cover glass (Warner Instruments, number 0 cover glass, 3 mm in diameter; Norland Optics, number 81 adhesive). Any excess glass extending beyond the cannula edge was removed with a diamond-tipped file. During the imaging cannula implantation procedure, animals were anesthetized through intraperitoneal injection of ketamine (∼85 mg/kg) and xylazine (∼8.5 mg/kg). Anesthesia was maintained during the procedure with inhaled 0.5–1.5% isoflurane and oxygen at a flow rate of 0.8-1 L/min using a standard isoflurane vaporizer. Prior to surgery, animals received a subcutaneous administration of ∼2 mg/kg dexamethasone and 5-10 mg/kg Rimadyl (to reduce inflammation and promote analgesia, respectively). First, the viral injection site targeting the left dorsal CA1 was determined by stereotaxis (AP -1.94 mm, ML -1.10 to -1.30 mm) and an initial hole was drilled only deep enough to expose the dura. An automated Hamilton syringe microinjector (World Precisions Instruments) was used to lower the syringe (35-gauge needle) to the target depth at the CA1 pyramidal layer (DV -1.33 to -1.37 mm) and inject 500 nL adenovirus (AAV) at 50 nL/min, to express the genetically encoded calcium indicator GCaMP under the pan-neuronal synapsin promoter (AAV1-Syn-jGCaMP7f- WPRE, AddGene, viral prep 104488-AAV1, titer 2x10^12^). The needle was left in place for 10 min to allow for virus diffusion.

After retracting the needle, a 3 mm diameter circular craniotomy was then performed over the left hippocampus using a robotic surgery drill for precision (Neurostar). The craniotomy was centered at AP -1.95 mm, ML -1.8 to -2.1 mm (avoiding the midline suture). During drilling, the skull was kept moist by applying cold sterile artificial cerebrospinal fluid (ACSF; sterilized using a vacuum filter). The dura was then delicately removed using a bent 30-gauge needle. To access CA1, the cortex overlying hippocampus was carefully aspirated using a blunt aspiration needle, with continuous irrigation of ice-cold, sterile ACSF. Aspiration was stopped when the fibers of the external capsule were clearly visible, leaving the external capsule intact. Following hemostasis, the imaging cannula was gently lowered into the craniotomy until the cover glass lightly contacted the fibers of the external capsule. To optimize an imaging plane tangential to the CA1 pyramidal layer while minimizing structural distortion, the cannula was positioned at an approximate 10° roll angle relative to horizontal. The cannula was affixed in place using cyanoacrylate adhesive. A thin layer of adhesive was also applied to the exposed skull surface, which was pre-scored with a number 11 scalpel before the craniotomy to provide increased surface area for adhesive binding. A headplate featuring a left-offset 7-mm diameter beveled window and lateral screw holes for attachment to the imaging rig was positioned over the imaging cannula at a matching 10° angle. The headplate was cemented to the skull using Metabond dental acrylic dyed black with India ink or black acrylic powder.

Upon completion of the procedure, animals were administered 1 mL of saline and 10 mg/kg of Baytril, then placed on a warming blanket for recovery. Monitoring continued for the next several days, and additional Rimadyl and Baytril were administered if signs of discomfort or infection appeared. A minimum recovery period of 10 days was required before initiation of water restriction, head-fixation, and virtual reality training.

### Histology

After the conclusion of all experiments, mice were deeply anesthetized and administered an overdose of Euthasol, then perfused transcardially with PBS followed by 4% paraformaldehyde (PFA) in 0.1M PB. Brains were removed and post-fixed in PFA for 24 hours, followed by incubation in 30% sucrose in PBS for >4 days. 50 µm coronal sections were cut on a cryostat, mounted on gelatin-coated slides, and coverslipped with DAPI mounting medium (Vectashield). Histological images were taken on a Zeiss widefield fluorescence microscope.

### Virtual Reality (VR) Design

All VR tasks were designed and operated using custom code written for the Unity game engine (https://unity.com). Virtual environments were displayed on three 24-inch LCD monitors surrounding the mouse at 90° angles relative to each other. The VR behavior system included a rotating fixed axis cylinder to serve as the animal’s treadmill and a rotary encoder (Yumo) to read axis rotations to record the animal’s running. A capacitive lick port, consisting of a feeding tube (Kent Scientific) wired to a capacitive sensor, detected licks and delivered sucrose water reward via a gravity solenoid valve (Cole Palmer). Two separate Arduino Uno microcontrollers operated the rotary encoder and lick detection system. Unity was controlled on a separate computer from the calcium imaging acquisition computer. Behavioral data were sampled at 50 Hz, matching the VR frame rate. Both the start of the VR task as well as each 50 Hz frame were synchronized with the ∼15.5 Hz sampled imaging data via Unity-generated TTL pulses from an Arduino to the imaging computer.

### Behavioral training and VR tasks

#### Handling and pre-training

After ∼1 week of recovery, the mice were handled for 2-3 days for 10 minutes each day, then acclimated to head-fixation (at least 10 days after surgery) on the cylindrical treadmill in a stepwise fashion for at minimum 15-min sessions for 2-3 days (increasing to 30 min-1 hour on the second to third day). Mice were then acclimated to the lick port by 1-2 days of pre-training to lick for water rewards delivered at 2- second intervals (i.e. a minimum of 2 s between rewards if mice were actively licking, otherwise no reward was delivered). For this pre-training and all subsequent behavior, sucrose water reward (5% sucrose w/v) was delivered when the mouse licked the capacitive lick port. The water delivery system was calibrated to release ∼4 μL of liquid per drop. To motivate behavior, mice were water restricted to 85% or higher of their baseline body weight. Mice were weighed daily to monitor health and underwent hydration assessments (via skin tenting). The total volume of water consumed during the training and behavioral sessions was measured, and after the experiment each day, supplemental water was supplied up to a total of ∼0.045 mL/gm per day (typically 0.8-1 mL per day, adjusted to maintain body weight). After the training and water consumption, the animals were returned to their home cage each day.

Once acclimated to the lick port, mice were pre-trained on a “running” task on a 350 cm virtual linear track with random black and white checkerboard walls and a white floor to collect a cued reward. The reward location was marked by two large gray towers, initially positioned at 50 cm down the track. If the mouse licked within 25 cm of the towers, it received a liquid reward; otherwise, an automatic reward was given at the towers. After dispensing the reward, the towers advanced (disappeared and quickly reappeared further down the track). Covering the distance to the current reward in under 20 s increased the inter-reward distance by 10 cm, but if the mouse took longer than 30 s, the distance decreased by 10 cm, with a maximum reward location of 300 cm. Upon consistent completion of 300 cm distance in under 20 s, the automatic reward was turned off, requiring the mice to lick to receive reward. At the end of the track, the mice passed through a black “curtain” and into a gray teleport zone (50 cm long) that was equiluminant with the VR environment before re-entering the track from the beginning. Once mice were reliably licking and completing 200 laps of the pre-training track within a 30-40 min period, they were advanced to the main task and imaging (mean ± s.d.: 6.1 ± 1.8 days of running pre-training across 10 mice).

#### Hidden reward zone “switch” VR task

The main “switch” task reported in this study involved two virtual environments highly similar to those previously used to study hippocampal remapping^7^, each with visually distinct features from the pre-training environment. Both environments consisted of a 450 cm linear track, with two colored towers and two patterned towers along the walls. Environment 1 (Env 1) had diagonal low-frequency black and white gratings on the walls, a gray floor and dark gray sky, and began with two green towers. Environment 2 (Env 2) had higher frequency gratings on the wall, a gray floor, a very light gray sky, and began with two blue towers. The reward zone was a “hidden” (not explicitly marked) 50-cm span at one of 3 possible locations along the track, each equidistantly spaced between the towers but with different proximity to the start or end of the track: zone A: 80-130 cm, zone B: 200-250 cm, zone C: 320-370 cm. Only one reward zone was ever active at a time. On the first 10 trials of any new condition, including the first day and each switch subsequently described, the reward was automatically delivered at the end of the zone if the mouse had not yet licked within the zone to signal reward availability. Otherwise, reward was delivered at the first location within the zone where the mouse licked. After these 10 trials, reward was operantly delivered for licking within the zone. Reward was randomly omitted on approximately 15% of trials (i.e. if a random number generator exceeded 0.85). Each lap terminated in a black curtain and gray teleportation “tunnel” to return the mouse to the beginning of the track. Time in the teleport tunnel was randomly jittered between 1 and 5s (5-10s on trials following reward omissions or trials where the mouse did not lick in the reward zone), followed by 50 cm of running during which the beginning of the track was visible to provide smooth optic flow.

Each mouse encountered a different starting reward zone and sequence of reward zone switches, counterbalanced across mice (n = 7 mice). Mice were allowed to learn an initial reward zone (e.g. A as in **Fig. 1E**) for days 1-2 of the task. On day 3 (Switch 1), the zone was moved to one of the two other possible locations on the track after 30 trials (e.g. B). For the first 10 trials at the new location, automatic delivery was activated at stated above, but empirically, we observed that the mice often started licking at the new location before that 10 trials had elapsed (**Fig. S1A-C**). The new reward zone was maintained on day 4. On day 5 (Switch 2), the zone was moved to the third possible reward zone (e.g. C), maintained on day 6, and moved back to the original location on day 7 (e.g. A). Each switch occurred after 30 trials. On day 8, the reward zone switch coincided with a switch into Env 2, where the sequence of zone switches was then reversed on the same day-to-day schedule for a total of 14 days (**Fig. 1**, **Fig. S1**).

An additional “fixed-condition” cohort (n=3 mice) experienced only Env 1 and one fixed reward location throughout the 14 days (**Fig. S1D,F**).

We targeted 80-100 trials per session with simultaneous two-photon imaging (described below), with 1 imaging session per day (mean ± s.d: 80.9 ± 8.8 trials across 10 mice). The session was terminated early if the mouse ceased licking for an extended period of time or ceased running consistently, and/or if the imaging time exceeded 50 minutes. Prior to the imaging session for this task, mice were provided 30 “warm-up” trials using the task and reward zone from the previous day to re-acclimate them to the VR setup. On day 1 of imaging, this warm-up session used the pre-training environment from prior days. On days 2 onward, the warm-up session was whichever environment and reward zone was active at the end of the previous day. Following the completion of the daily imaging session, mice were given another ∼100 training trials without imaging on the last reward zone seen in the imaging session until they acquired their daily water allotment.

### Two-photon (2P) imaging

We used a resonant-galvo scanning 2P microscope from Neurolabware to image the calcium activity of CA1 neurons. Excitation was achieved using 920-nm light from a tunable output femtosecond pulsed laser (Coherent Chameleon Discovery). Laser power modulation was accomplished with a Pockels cell (Conoptics). The typical excitation power, measured at the front face of the objective (Leica 25X, 1.0NA, 2.6 mm working distance), ranged from 13-40 mW for mice m12, m13, m14; 15-68 mW for mice m2, m3, m4, m6, m7; and 50-100 mW for m10 and m11. The hidden reward zone task was imaged starting from day 1 for all mice except m11, for whom imaging started on day 3 due to lower viral expression. Continuous imaging under constant laser power persisted through each trial of the imaging session. For most animals and sessions, to minimize photodamage and photobleaching, the Pockels cell was used to reduce laser power to minimal levels between trials (during the teleport zone). For mice m11, m12, m13, and m14, laser power was maintained and imaging continued throughout the teleport zone on days 1, 3, 7, 8, and 14 of the task. Photons were collected using Hamamatsu gated GAsP photomultiplier tubes (part H11706-401 for Neurolabware microscopes). Imaging data were acquired via Scanbox software (https://scanbox.org/), operated through MATLAB (Mathworks). The imaging field of view (FOV) was collected at 1.0 magnification with unidirectional scanning at ∼15.5 Hz, resulting in a ∼0.64 x 0.64 mm FOV (512 x 716 pixels). The same FOV was acquired each session by aligning to a reference image from previous days prior to the start of data acquisition, with the aid of the ‘searchref’ plugin in Scanbox. This allowed us to track single cells across days (see *Calcium data processing*).

### Calcium data processing

The Suite2P software package^174^ (v0.10.3) was used to perform xy motion correction (rigid and non-rigid) and identify putative cell regions of interest (ROIs). Manual curation was performed to eliminate ROIs containing multiple somas or dendrites, lacking visually obvious transients, suspected of overexpressing the calcium indicator, or exhibiting high and continuous fluorescence fluctuation typical of putative interneurons. This approach yielded between 155 and 1982 putative pyramidal neurons per session, due to variation in imaging window implant quality and viral expression. Custom code was used to identify cells that were successfully tracked across imaging sessions using the intersection over union (IOU) of their ROI pixels. The threshold for ROI matching was chosen algorithmically for each dataset such that the IOU for the best match for an ROI pair was always greater than the IOU for any second best match.

To compute the ΔF/F (dF/F) for each ROI, baseline fluorescence was calculated within each trial independently using a maximin procedure with a 20-s sliding window (modification of default suite2p procedure). Limitation to individual trials both accounts for potential photobleaching over the session and avoids the teleport periods for sessions where the laser power was reduced (see *Two-photon imaging*). dF/F was then calculated for each cell as the fluorescence minus the baseline, divided by the absolute value of the baseline. dF/F was smoothed with a 2-sample (∼0.129 s) s.d. Gaussian kernel. The activity rate was then extracted by deconvolving dF/F with a canonical calcium kernel using the OASIS algorithm^175^ as used in suite2p. It is important to note that this deconvolution is not interpreted as a spike rate but rather as a method to eliminate the asymmetric smoothing effect on the calcium signal induced by the indicator kinetics. Additional putative interneurons were detected for exclusion from further analysis via a Pearson correlation >0.5 between their dF/F timeseries and the animal’s running speed, excluding 0.24 ± 0.41% of cells (mean ± standard deviation across mice and days).

### Statistics

To avoid assumptions about data distributions, we used nonparametric tests or permutation tests in most cases, except where a Shapiro–Wilk test for normality confirmed that parametric tests were reasonable. Specific tests are described in further detail in specific Methods sections. In the main text, percentages of cells classified by remapping type are reported as mean ± standard deviation out of all place cells identified on the specified days (n = 7 mice for the “switch” task). Place cell criteria are defined below in *Place cell and peak firing identification*. Averaged data in figures are shown as mean ± s.e.m. unless otherwise indicated. For the distributions of sequence positions over days in Fig. 3 and Fig. S3, s.e.m. shading is omitted for clarity, but the variance for these distributions is taken into account in linear mixed effects models to quantify these plots (see *Linear mixed effects models*).

All analyses were performed in Python 3.8.5. Linear mixed effects models were performed using the statsmodels package (https://www.statsmodels.org/stable/index.html). Pairwise repeated measures (paired) t-tests were run using the pingouin package (https://pingouin-stats.org/) with a Holm step-down Bonferroni correction for multiple comparisons. Linear regressions and Wilcoxon signed-rank tests were performed using the SciPy (https://scipy.org/) statistics module, with confidence intervals computed with the Uncertainties package (https://pypi.org/project/uncertainties/). Circular statistics were performed using Astropy (https://docs.astropy.org/en/stable/stats/circ.html), pycircstat^176^ (https://github.com/circstat/pycircstat), and circular-circular correlation code originally written to analyze hippocampal theta phase precession^177^ (https://github.com/CINPLA/phase-precession). The time warp models and GLM were implemented from publicly available repositories (time warp: https://github.com/ahwillia/affinewarp; GLM: https://github.com/HarveyLab/GLM_Tensorflow_2).

The number of mice to include was determined by having coverage of every possible reward sequence permutation (6 sequences) with at least one mouse. In addition, our mouse sample sizes were similar to those reported in previous publications^7,150^.

### Quantification of licking behavior

The capacitive lick sensor allowed us to detect single licks. A very small number of trials where the sensor had become corroded were removed from subsequent licking analysis by setting these values to NaN (∼0.8% of all imaged trials, n=64 removed trials out of 7736 trials across the 7 switch mice). These trials were detected by >30% of the imaging frame samples in the trial containing ≥3 Unity frames where licks were detected, as this would have produced a sustained lick rate ≥20 Hz. Remaining lick counts were converted to a binary vector at the imaging frame rate and spatially binned at the same resolution as neural activity (10 cm bins), then divided by the time occupancy in each bin to yield a lick rate. We quantified the licking precision of the animals over blocks of 10 trials using an anticipatory lick ratio inspired by the ramp of anticipatory licking we observed preceding the reward zone (**Fig. 1F-G, Fig. S1A-C**). The ratio was computed as:

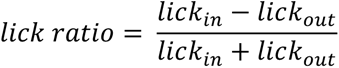

Where *lick_in_* is the mean lick rate in a 50 cm “anticipatory” zone before the start of the reward zone, and *lick_out_* is the mean rate outside of this zone and the reward zone. The reward zone itself is thus excluded to exclude consummatory licks (**Fig. S1E**). A ratio of 1 indicates perfect licking only in the anticipatory zone, a ratio of -1 indicates licking only outside of this zone, and a ratio of 0 indicates chance licking everywhere excluding the reward zone.

### Place cell and peak firing identification

For all neural spatial activity analyses we excluded activity when the animal was moving at <2 cm/s. To get a trial-by-trial activity rate matrix for each cell, we binned the 450 cm linear track into 45 bins of 10 cm each. For each cell, we took the mean of the activity samples (dF/F or deconvolved calcium activity) on each trial within each position bin (note that taking the mean activity over time samples is equivalent to normalizing by the occupancy within a trial), producing a matrix of trials x position bins.

We defined place cells as cells with significant spatial information^178^ (SI) in either trial set 1 (pre-switch) *or* trial set 2 (post-switch) in a session. On days without a switch, we used the 1st and 2nd half of trials. For the spatial information computation and for plotting spatial firing over trials, we used the deconvolved calcium activity as this reduces the asymmetry of the calcium signal (smoothed only for plotting with a 10 cm s.d. Gaussian kernel). SI was calculated using a formula implemented for calcium imaging^179^ as:

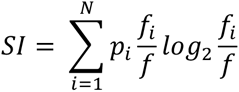

where *p_i_* is the occupancy probability in position bin *i* for the whole session, *f_i_* is the trial-averaged activity per position bin *i*, and *f* is the mean activity rate over the whole session, computed as the sum of *f_i_ * p_i_* over all *N* bins. To get *p_i_,* we calculated the occupancy (number of imaging samples) in each bin on each trial and divided this by the total number of samples in each trial to get an occupancy probability per position bin per trial. We then summed the occupancy probabilities across trials and divided by the total per session to get an occupancy probability per position bin per session. To get the spatial “tuning curve” over the session, we averaged the activity in each bin across trials. To determine the significance of the SI scores, we created a null distribution by circularly permuting the position data relative to the timeseries of each cell, by a random amount of at least 1 s and a maximum amount of the length of the trial, independently on each trial. SI was calculated from the trial-averaged activity of each shuffle, and this shuffle procedure was repeated 100 times per cell. A cell’s true SI was considered significant if it exceeded 95% of the SI scores from all shuffles within animal (i.e. shuffled scores were pooled across cells within animal to produce this threshold, which is more stringent than comparing to the shuffle of each individual cell^180^).

To identify spatial peaks, we used the unsmoothed spatially binned dF/F, as this signal is the closest to the raw data. Place cell firing peaks or “spatial peaks” refer to the position bin of the maximum spatially-binned activity, usually averaged across trials within a set. There was no restriction to field boundaries, thus allowing cells to have multiple fields. To plot single cell place fields and cell sequences, neural activity was normalized to the mean of each cell within a session.

### Trial-by-trial similarity matrices

Correlation matrices for single cells were computed using the spatially binned deconvolved activity on each trial, smoothed with a 2-bin (20 cm) s.d. Gaussian. This resulted in a matrix *A_i_* for each cell *i* of *J* trials x *M* position bins. Each trial was z-scored across the position axis, and the trial-by-trial correlation matrix *C* was computed as:

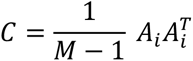

### Cell category definitions for remapping subtypes

We defined the remapping types shown in **Fig. 2** and **Fig. S2** as follows: *track-relative*: significant SI before and after the switch, with spatial peaks (position bin of maximum trial-averaged dF/F) ≤50 cm from each other before vs. after; *disappearing*: significant SI before but not after the switch, with a mean spatially-binned dF/F after that is less than the 50th percentile of the per-trial mean dF/F before; *appearing*: significant SI after but not before, with a mean spatially-binned dF/F after that is greater than 1 s.d. above the mean in trials before; *remap near reward*: significant SI before and after, with spatial peaks ≤50 cm from the starts of both reward zones; *remap far from reward*: significant SI before and after, not “track-relative” (i.e. peaks >50 cm apart before vs. after) and not “near reward” cells (i.e. peaks >50 cm from the start of at least one reward zone).

### Reward-relative remapping

#### Quantification of reward-relative remapping compared to chance

To perform statistics on putative reward-relative remapping (**Fig. 2, Fig. S2**), we restricted the set of included place cells to require significant spatial information in the 1st *and* 2nd trial set in each session (pre- and post- switch), to produce a stringent set of spatial peaks. For all scatter plots of spatial peaks before vs. after a reward switch, points are jittered by a random amount between -π /100 and +π/100 for visualization only. As outlined in **Fig. 2**, we converted the track axis (i.e. the original position timeseries of the animal) to circular coordinates, setting the beginning of the track to -π and the end of the track (beginning of the teleport zone) to π. Spatial peaks for each cell were re-computed using dF/F binned into 45 bins on this axis (bin size 2π/45, corresponding to ∼10 cm). Putative track-relative cells were removed by excluding cells with spatial peaks before vs. after the switch that fell within 0.698 radians (∼50 cm) of each other. The coordinates of the remaining data were then circularly rotated to align the start of each reward zone at 0, such that spatial peaks could be measured relative to the start of each reward zone (as the signed circular distance). We measured the distance between relative peaks as the “after” peak minus the “before” peak, circularly wrapped, creating a distribution that can be thought of as orthogonal to the unity line shown in the scatters in **Fig. 2D, F**. We compared this distribution (shown in **Fig. 2E, G**) to a “random-remapping” shuffle. This shuffle was generated by maintaining the pre-switch position of each cell’s peak and circularly permuting the cell’s post-switch firing 1000 times by a random offset of 0 to 44 bins, creating a set of shuffled post-switch positions for each cell. To define a candidate range of reward-relative remapping variability (i.e. a range around the unity line in **Fig. 2D, F**), we computed the circular difference between the maximum and minimum bin of the distribution that exceeded the upper 95% of the shuffle, divided by 2, and averaged across switches (**Fig. S2G**). This produced a mean range of ∼0.656 radians (46.9 cm) around 0, which is captured by a 50 cm threshold given our 10 cm bin size. This threshold was subsequently used to identify candidate reward-relative remapping cells (see below).

#### Criteria to define reward-relative cells

Reward-relative cells could encompass “remap near reward” and “remap far from reward” cells as defined in *Cell category definitions for remapping subtypes*. However, we observed variable dynamics over trials with which the spatial activity of these cells remapped after a reward switch (**Fig. 2A-B, Fig. S2A**), which could affect spatial information scores. We therefore relaxed the SI criterion for reward-relative cells such that they were required to have significant SI either before or after the switch but not necessarily both, to allow for these dynamics. We began by finding place cells which had peak spatial firing within ≤50 cm (0.698 radians) of each other when these peaks were calculated using circular bins relative to reward zone starts. As the detection of peak firing can be noisy, we further took into account the shape of the field by performing a cross-correlation between each place cell’s trial-averaged spatial activity before vs. after, using the activity matrices (trials x position bins, unsmoothed dF/F) where reward zone starts are circularly aligned between trial sets 1 and 2. This procedure allowed us to test the hypothesis that reward-relative cells should have a peak cross-correlation close to zero on this reward-centered axis. We calculated a shuffle for the cross-correlation by circularly permuting the activity in trial set 2 by a random offset between 1 and 45 bins on each trial 500 times, and required that the real cross-correlogram had a peak exceeding the 95% confidence interval (upper 97.5% threshold) of the shuffle, and that the offset of this peak was within 5 bins (∼50 cm) of zero (see **Fig. S2N-P**). We validated that the population of cells passing this criterion had high Pearson correlation coefficients between their circularly aligned activity maps in set 1 vs. set 2 (whereas, by contrast “track-relative” cells had high correlation coefficients and a cross-correlation offset near zero only in the original linear track coordinates; **Fig. S2O-P**). Place cells that passed this final circular cross-correlation criterion were therefore defined as reward-relative.

### Relationship between distance run in the teleport zone and trial-to-trial variability

We measured the distance run in the teleport zone at the end of each trial by integrating the animal’s speed between the entry to the teleport zone (i.e. first frame at which the temporal jitter started, see *Behavioral training and VR tasks*) and the last frame before the re-entry to the track (i.e. start of the next trial). We then computed the trial-wise spatial peak error for each reward-relative cell as phase difference between the cell’s spatial peak location in circular coordinates on each trial and its mean spatial peak location within either trial set 1 (before the reward switch) or trial set 2 (after the switch). Circular error was then converted back to centimeters. We next performed a Spearman correlation between distance run during the teleport and the error on the following trial; population correlation coefficients are reported in **Fig. S2F**. Results were similar using linear root-squared error rather than circular coordinates (data not shown).

### Sequence detection and quantification

The sequential firing of neural subpopulations was detected using a split-halves cross-validation procedure^7^. For these analyses, we used the unsmoothed, spatially binned dF/F in circular track coordinates (not aligned to reward, though aligning to reward makes no difference for the circular sequence order). To sort the activity of neurons, we used the activity averaged over the odd trials before the switch to find and sort the peak firing positions. This sort was validated by plotting the trial-averaged activity of the even trials before the switch (e.g. left-hand columns of **Fig. 3A**). The same sort was then applied to the trials after the switch. The sequence positions before vs. after were then taken as the peak firing positions of the trial-averaged “even-before” trials and the trial-averaged “after” trials. Preservation of sequence order was quantified as the circular-circular correlation coefficient^176,177,181^ between the sequence positions before vs. after. Though a p-value can be directly computed for this correlation coefficient, its significance was further validated through a permutation procedure. We randomly permuted the cell identities of the neurons after the switch and re-computed the correlation coefficient 1000 times to get a null distribution. The observed correlation coefficient was considered significant if it was outside 95% of the null distribution using a two-tailed test. The p-value was calculated as:

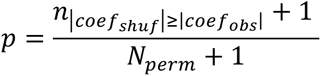

where n|*coefshuf*|≥|*coefobs*| is the number of shuffled coefficients with absolute values ≥ the absolute value of the observed coefficient, and *N_perm_* is the total number of shuffles.

For sequences tracked across days, we again used the cross-validated sort from trial set 1 (before the switch, or the 1st half of trials if the reward did not move) on the reference day (the first day in each day pair) and applied this sort to trial set 2 (after the switch, or 2nd half) on the reference day as well as activity in trial sets 1 and 2 on the target day (the second day in the pair). To measure drift between days, we computed the across-day circular-circular correlation coefficient between the sequence positions on reference day trial set 2 vs. target day trial set 1.

To quantify the density of sequences, we divided the number of cells in the sequence with peaks in each position bin by the total number of place cells for that animal. These density curves were smoothed only for visualization in **Fig. 3J, S** and **Fig. S3J** with a 1-bin (2π/45) s.d. Gaussian and shown as the mean across animals per each switch day. To visualize the expected uniform distribution across animals, we calculated this as the mean fraction of cells in each sequence out of the place cells in each animal on a given switch day, divided by the number of position bins.

### Linear mixed effects models

To quantify neural activity changes across task experience, we used linear mixed effects models to take into account variance across animals. Linear mixed effects models were implemented using the mixedlm method of the statsmodels.formula.api package (https://www.statsmodels.org/stable/mixed_linear.html). Reported p-values are from Wald tests supplied by mixedlm. The fixed effect was either the switch index or day-pair index as a continuous variable, except for **Fig. S5J** where additional fixed effects were considered (see figure legend). Random effects were mouse identity. Random intercepts for each mouse were allowed; we also confirmed that including random slopes did not affect the results. When the dependent variable was reported in fractions of cells, we applied a logit transform to the fractions when a large proportion of the values were near zero. In most cases, however, we chose to display results in the original fractions for interpretability. In these cases, we confirmed that performing a logit transform on the fractions did not qualitatively change the results and only modestly changed the model fit.

### Analysis of rewarded vs. omission trials

All following analyses were restricted to trial sets (i.e. before or after the reward switch) that had at least 3 omission trials within the set.

#### Time warp modeling

We first fit 5 different time warp model types—shift, linear, piecewise 1 knot, piecewise 2 knots, and piecewise 3 knots^87^—on the matrix of speed profiles within a trial set (T trials x M linear position bins x 1, expanded in the third dimension for compatibility with the time warp algorithm). These models apply warping functions of increasing nonlinear complexity to stretch and compress the data on each trial for maximal alignment^87^. We included both rewarded and omission trials in the fitting procedure to find the best alignment across them. Model fit was assessed using the mean squared error (MSE) between the time warped speed profile of each trial and the mean across time warped trials (**Fig. S5A**). The piecewise 3 knots model most often produced the lowest MSE, or best fit, across sessions (**Fig. S5C**). We therefore re-fit all sessions with piecewise 3 knots to ensure that model selection could not influence the reward vs. omission metric described below. We then applied the model transform to the neural data matrix for the same set of trials (T trials x M linear position bins x N neurons; excluding neural data at movement speeds <2cm/s, similar to all other neural analyses). For neural analysis in **Fig. 5**, we then focused on trial sets before the switch where the reward zone was at position “A” or “B”. This is because the mean rewarded vs. omission speed profiles were more similar on these trials than after the switch (**Fig. S5F**) or when the reward was at “C” (as the animals tended to lick for longer on omission trials and there is less room at the end of the track for running speed to reach its maximum from position “C”; see **Fig. S1** and **Fig. S5H** for examples). Restricting sessions to “A” and “B” reward zones produced similar results to restricting sessions to those best fit by the time warp model (data not shown). Complementary analyses for trials after the switch and for all reward zones are reported in **Fig. S5**.

#### Reward vs. omission index

We compared neural activity on rewarded vs. omission trials for reward-relative cells with spatial peaks between the start of the reward zone and the end of the track (under the hypothesis that these cells could receive information about whether reward was received or not on that trial). Using the time warp model-transformed neural activity averaged across rewarded or omission trials within the trial set of interest, we calculated a “reward vs. omission index” (RO index) for each of these reward-relative cells as:

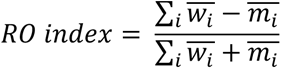

where *w̄_ī_* is the mean activity in position bin *i* averaged across rewarded trials, and *m̄_ī_* is the mean activity in position bin *i* averaged across omission trials. RO indices of 1 therefore indicate an exclusive firing preference for rewarded trials, -1 indicates an exclusive preference for omission trials, and 0 indicates equal firing between rewards and omissions. We confirmed that the RO index calculated from the model- transformed activity was highly correlated with the RO index calculated from the original activity (**Fig. S5E**). We then quantified the median RO index for each animal and day. Using a linear mixed effects model (**Fig. SJ**), we found that there was no significant change across days, as a result of the model fit, or as a result of the original MSE between rewarded and omission trials. We therefore combined neurons across days for presentation in **Fig. 5E**.

### Generalized linear model (GLM)

We implemented a Poisson GLM developed previously^88^ to predict the deconvolved calcium event time series of individual neurons from a set of task and movement variables. All behavioral and neural time series were sampled at ∼15.5 Hz, the imaging frame rate.

#### Design matrix

Our design matrix is schematized in **Fig. S6A** and included 6 predictor variables split into 3 task variables and 3 movement variables.

Task variables were linear track position (from 0 to 450 cm), reward-relative position (from -π to π, centered around the start of each reward zone at 0 radians), and “rewarded”, a binary that goes from 0 to 1 when the reward is delivered on each trial and stays at 1 until the end of the trial (rewarded stays at 0 on omission trials). Linear track position and reward-relative position were separately expanded into 45 cosine basis functions (or “bumps”) each, one for each 10 cm or ∼0.14 radian bin used in other analyses. The rewarded variable was multiple with the linear position basis functions to yield the interaction between position and whether the animal had been rewarded on a given trial.

Movement variables were running speed and acceleration (smoothed with a 5-sample s.d. Gaussian) and smoothed lick count (binary licks per frame smoothed with a 2-sample s.d. Gaussian to approximate an instantaneous rate). Each movement variable was separately quantile-transformed to encode the distribution of movement dynamics similarly across animals. The quantiles were expanded with B-spline basis functions using the SplineTranformer method of Python’s sklearn package, with 3 polynomial degrees and 5 knots, resulting in 7 bases (5+3-1) per movement variable. Spline choices were made for consistency with previous work^88^.

All predictors were concatenated into a design matrix with 156 total features: 45 position bases + 45 reward-relative position bases + 45 rewarded-x-position bases + 7 speed bases + 7 acceleration bases + 7 licking bases. The features were independently z-scored across samples before model fitting on the “response matrix” of deconvolved activity of all cells (samples x neurons).

#### Model fitting and testing

Trial identities were used to group data for training and testing. Individual sessions (including trials both before and after the reward switch) were split into 85% training trials and 15% testing trials, using the GroupShuffleSplit method of Python’s sklearn package with a consistent random seed. Specifically, trials were allocated such that the training trials included 85% of rewarded trials before the switch, 85% of rewarded trials after the switch, and 85% of all omission trials (likewise the testing set received the remaining trials from each group). The training trials were further divided for 5-fold cross validation during the fitting procedure. An optimal model was selected according to the deviance explained on the 5-fold cross-validation data. We then tested this model on the held-out test data to assess model performance as the fraction deviance explained (FDE):

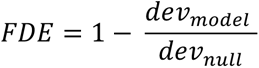

where *dev_model_* is the Poisson deviance of the full model and *dev_null_* is the Poisson deviance of a null model that predicts the mean of the neural activity across time samples.

On average, the model’s FDE for all place cells was 0.10 ± 0.19 (mean ± s.d.) (**Fig. S6B-C**), 0.32 ± 0.13 for track-relative cells, 0.29 ± 0.11 for reward-relative cells (examples in **Fig. 6A-B**), and 0.29 ± 0.11 for non-reward-relative remapping cells. For analysis of the relative contribution of individual variables, we only included cells with FDE > 0.15 in accordance with previous procedures^88^.

#### Relative contribution of individual variables via model ablation

To assess how much each variable contributed to the model’s ability to predict a given cell’s activity, we performed a model ablation procedure. In brief, after fitting the full model, we zeroed the coefficients for each variable and compared the performance of the ablated versus the full model on the cross-validation data. We calculated the reduction in model fit (deviance) of the ablated model relative to the full model, normalized by the full model’s deviance relative to the null model. This is known as “fraction explained deviance”^88^, which we term “relative contribution” to distinguish it from fraction deviance explained. Relative contribution was thus computed as:

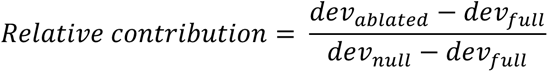

Relative contribution was binned by linear track position to visualize the contribution at each position along the track, then averaged across position bins for quantifying the contribution at the population level (**Fig. 6D**) and identifying the top predictor per cell (variable with the maximum mean relative contribution) (**Fig. 6E**).

---

**Figure S1.**
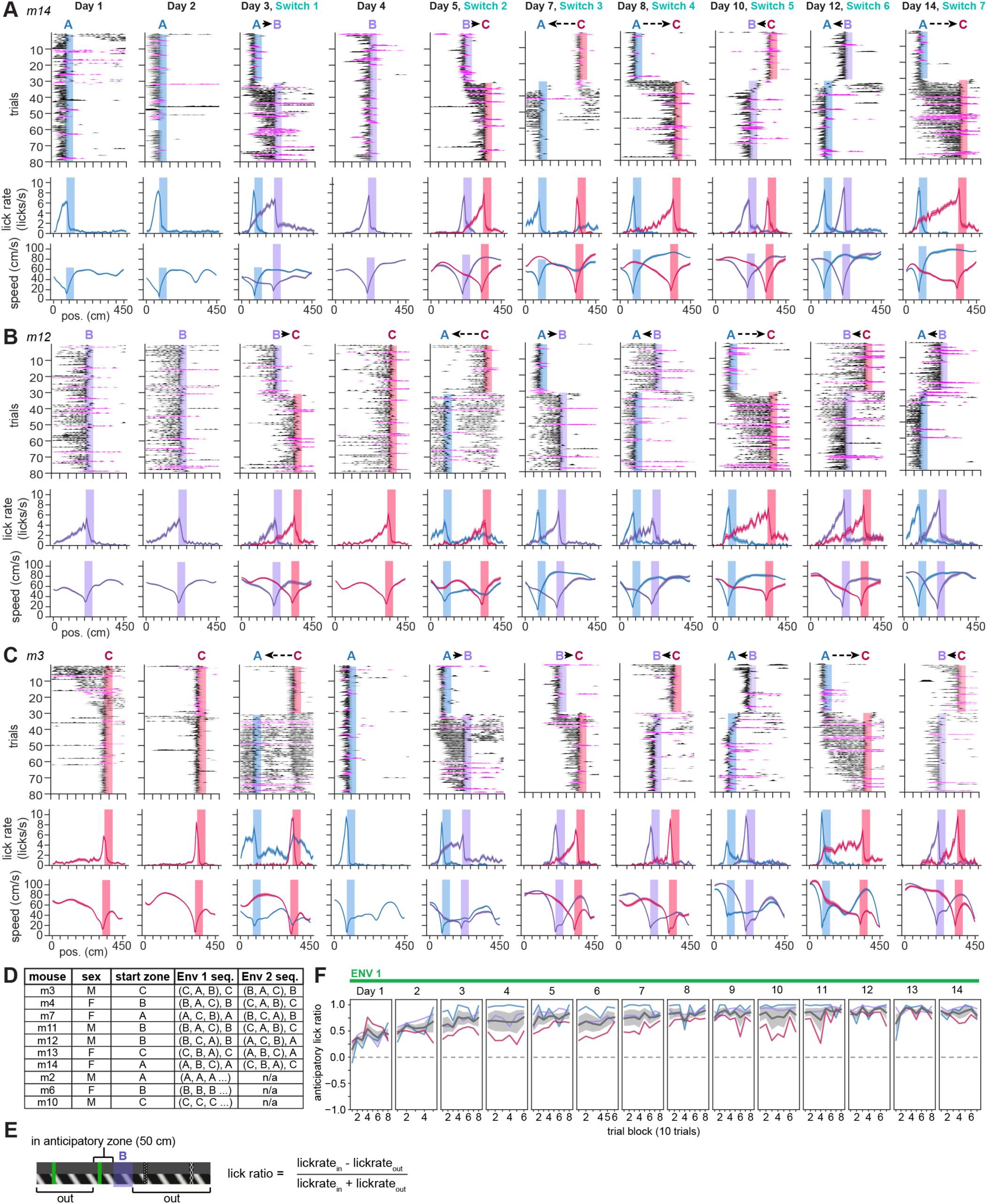
Behavior across multiple reward switch sequences, related to Figure 1. **A)** Expanded behavior over the 14-day task for the example animal in Fig. 1F which started the task with reward zone “A” (m14), showing days 1-5 and then each subsequent switch (day 7, 8, 10, 12, 14). Top: smoothed lick raster over 80 trials per session; black indicates rewarded trials, and magenta indicates omission trials. Shaded regions indicate the reward zone active on each set of trials. Middle: The mean ± s.e.m. lick rate for trials at each reward location (blue=A, purple=B, red=C). Bottom: Mean ± s.e.m. speed for trials at each reward location. **B)** Behavior for an example animal that started the task with reward zone “B” (m12), organized as in (A). **C)** Expanded behavior for the example animal in Fig. 1G which started the task with reward zone “C” (m3), organized as in (A). **D)** Mouse identities, sexes, and reward switch sequence allocation. Top 7 mice are “switch” mice, bottom 3 mice are “fixed-condition” mice that maintained the same reward zone and environment throughout the experiment. Sequences are shown as “(0, 1, 2), 0” where 0 is the first reward zone experienced in a given environment, 1 is the first switch (2nd zone), 2 is the second switch (3rd zone), and the third switch which is a return to zone 0. **E)** Schematic depicting anticipatory lick ratio calculation around reward zone “B” as an example. The anticipatory lick ratio compares lick rate in an anticipatory zone 50 cm before the start of the reward zone vs. everywhere outside the reward zone (the reward zone is excluded to eliminate consummatory licks). A value of 1 indicates licking exclusively in the anticipatory zone, 0 is chance everywhere outside the anticipatory zone and the reward zone. **F)** Anticipatory lick ratio across blocks of 10 trials for fixed-condition animals (n=3 mice) which maintained the same reward zone and environment throughout the entire experiment. Colored lines represent individual mice and the reward zone active for each mouse (blue=A, purple=B, red=C); gray lines and shading show mean ± s.e.m across mice.

**Figure S2.**
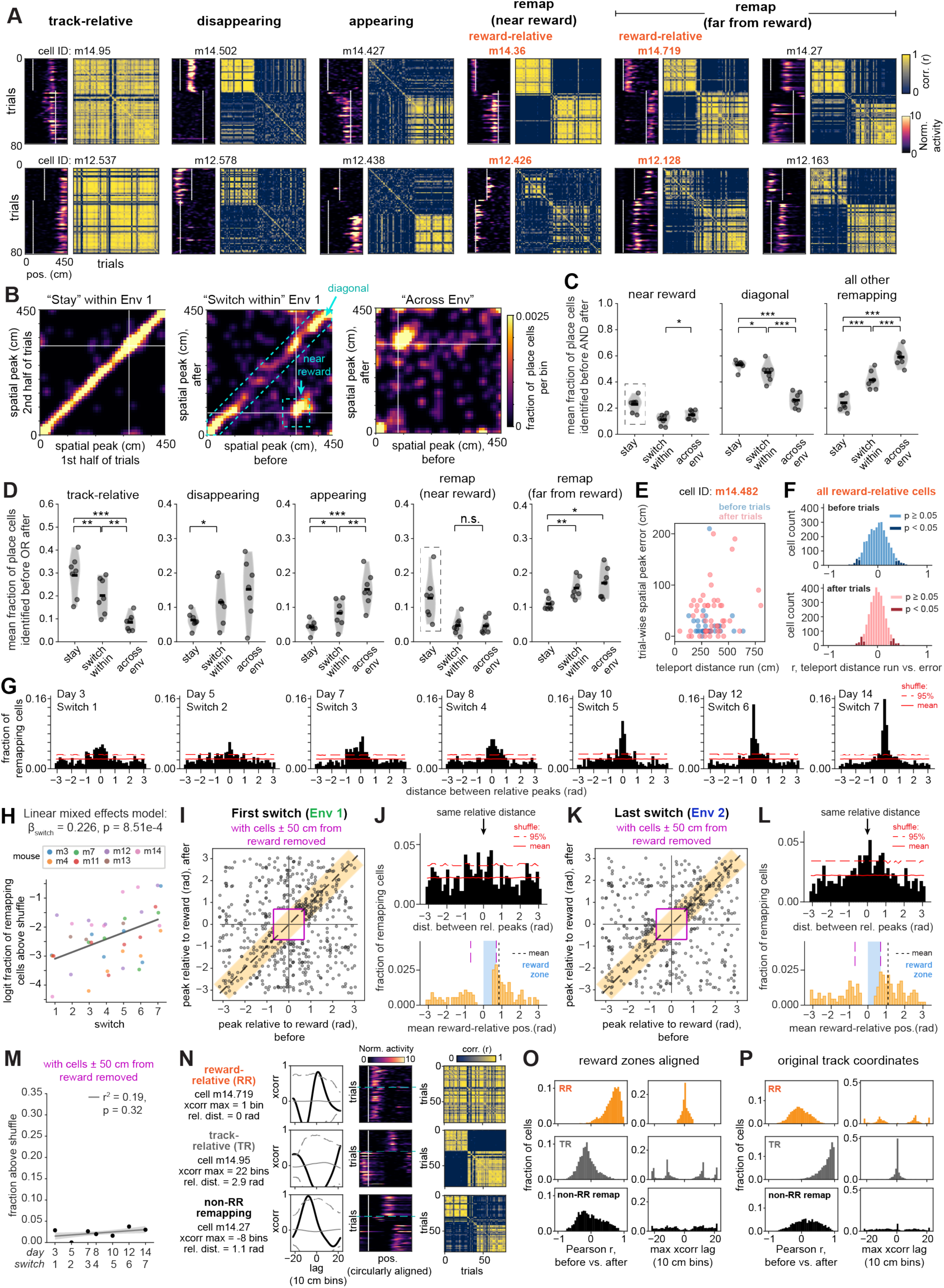
Quantification of remapping types and controls for reward-relative remapping, related to Figure 2. **A)** Example cells from 2 mice not shown in Fig. 2A on the last switch (day 14), replicating the co-existence of remapping types within animal shown in Fig. 2A. Top row: Data from mouse m14. Bottom row: Data from mouse m12. The “reward-relative” designation (orange text) encompasses cells both near and far from reward that maintain their firing distance with respect to the reward zone (see Fig. 2B and Methods). **B)** Examples of population maps from mouse m14 illustrating a day that is “stay” (day 6, same reward location, same environment), “switch within” (day 7, change reward location, same environment), or “across environment” (day 8, change reward location, change environment). Each plot shows the smoothed 2D histogram of the fraction of place cells with peak firing in each 10 cm bin before vs. after the reward zone switch (or in the 1st vs. 2nd half of trials on “stay” days). White lines indicate the start of each reward zone. For this analysis, only cells with significant spatial information both before and after are included. On stay days, this produces a strong density along the diagonal, corresponding to place cells that remain track-relative throughout the session. On switch days, this diagonal degrades as more cells remap, and the off-diagonal band around the reward intersection appears (see also Fig. 2C). Upon the across-environment switch, the diagonal degrades further indicating more global remapping, but the reward-relative band is maintained. Cyan boxes mark bins used for the “diagonal” and “near reward” categories shown in (C). **C)** Quantification of population-level remapping using bins of the histograms illustrated in (B) (unsmoothed for quantification). Each dot is the mean fraction of cells across days of each category and environment per animal (n=7 mice); violins show the distribution across mice, horizontal lines show the mean across mice. In (C) and (D), *p<0.05, **p<0.01, ***p<0.001 from pairwise paired t-tests between categories (“stay”, “switch within” environment, or “across env”), performed on the logit-transformed fractions. P-values are Holm corrected for multiple comparisons within each plot. Left: Near reward; cells with peaks ≤50 cm from both reward zone starts; gray dashed box signifies that on stay days, this value reflects the fraction of total place cells near the reward across the 1st to 2nd half of the session, thus no t-tests are shown for this category as the reward is not moved. Switch within vs. across env: t = -3.0, p = 0.024. Middle: Diagonal; peaks within ≤50 cm of the same linear position before vs. after. Stay vs. switch within: t = 2.6, p = 0.042; stay vs. across env: t = 11.82, p = 6.6e-5; switch within vs. across env: t = 8.2, p = 3.6e-4. Right: All other remapping; all non-near-reward and non-diagonal bins. Stay vs. switch within: t = -7.7, p = 2.5e-4; stay vs. across env: t = -29.4, p = 3.1e-7; switch within vs. across env: t = -9.1, p = 2.0e- 4. **D)** Mean fractions of total place cells defined as track-relative (peaks ≤50 cm apart), disappearing, appearing, remap near reward (≤50 cm from both reward zone starts), or remap far from reward (>50 cm from at least one reward zone start), averaged across days of each type per mouse as in (C). See (C) for description of statistical tests. Track- relative: Stay vs. switch within: t = 5.7, p = 0.0012; stay vs. across env: t = 18.4, p = 5.0e-6; switch within vs. across env: t=6.6, p = 0.0011. Disappearing: Stay vs. switch within: t = -3.6, p = 0.033; stay vs. across env: t = -2.5, p = 0.093; switch within vs. across env: t = -0.96, p = 0.37. Appearing: Stay vs. switch within: t = -3.2, p = 0.018; stay vs. across env: t = -8.6, p = 4.1e-4; switch within vs. across env: t = -5.9, p = 0.002. Remap near reward: switch within vs. across env: t = -0.23, p = 0.83. As in (C), the gray dashed box indicates a fraction near reward on stay days that cannot be identified as “reward-specific”, as there is no reward switch; thus no statistical comparisons are shown for this category. Remap far from reward: Stay vs. switch within: t = -4.8, p = 0.0093; stay vs. across env: t = -3.8, p= 0.018; switch within vs. across env: t = -1.6, p = 0.16. **E)** To test the possibility that the putative reward-relative neurons encode distance run since the last reward, we leveraged the variable length teleport zone to ask whether distance run in the teleport zone predicted an offset (error) in the cell’s spatial firing on the subsequent trial (see Methods). Here, an example reward-relative cell is shown (cell m14.482 on day 8, switch 4 across environments; also shown in Fig. 2B). Each dot is a trial (blue = trials before the reward switch, pink = trials after the switch). This cell shows no significant Spearman correlation between the teleport distance run and spatial peak error for either set of trials (before: r = -0.10, p = 0.62; after: r = 0.19, p = 0.18). **F)** Quantification of Spearman correlation coefficients between teleport distance run and spatial peak error on the subsequent trial, for all reward-relative cells across all animals and switch days (n = 3104 cells, 7 mice, 7 switch days). Histograms are stacked; dark shades highlight cells with significant correlation coefficients (p < 0.05): 7.6% of reward-relative cells on before trials, 8.9% on after trials. Light shades highlight non-significant cells: 92.4% of reward-relative cells on before trials, 91.1% on after trials. **G)** Histograms of the circular distance between relative peaks after minus before for remapping place cells as in Fig. 2E, G, shown for all switch days (days 3 and 14 are repeated from Fig. 2). n=7 mice; day 3 n=677 cells, day 5 n=583 cells, day 7 n=726 cells, day 8 n=706 cells, day 10 n=508 cells, day 12 n=506 cells, day 14 n=707 cells. **H)** Increase in the fraction of cells remapping relative to reward above the shuffle over experience, at the level of individual animals. Each point is color-coded by mouse (n=7 mice) and represents the fraction of cells (logit-transformed) exceeding the “random-remapping” shuffle calculated within each individual mouse and switch day. Regression coefficient β and p-value are from a linear mixed effects model with switch index as the fixed effect and mice as random effects (random intercepts were allowed, and using random slopes did not change the outcome of the model; data not shown). Gray line shows the model prediction. **I-M)** Control for Fig. 2D-H, excluding cells with peaks within ± 50 cm (∼0.698 radians) from both reward zone starts (indicated by the magenta lines), intended to exclude putative reward-dedicated cells reported previously^54^. (I) n=557 cells, 7 mice. (K) n=466 cells, 7 mice. All points are jittered slightly for visualization (see Methods). The fraction of cells remapping relative to reward at distances greater than 50 cm from reward still exceeds the shuffle (J, top; L, top). (J, bottom) and (L, bottom): the distribution of mean reward-relative positions for the remaining cells shown in the orange shaded region around the unity line in (I) and (K), respectively. The means of these distributions are significantly non-zero: (J) non-zero mean = 0.848, 95% confidence interval [lower, upper] = [0.443, 1.253], circular mean test, n = 163 cells. (L) non-zero mean = 1.092, 95% confidence interval [lower, upper] = [0.573, 1.612], circular mean test, n = 167 cells. Black dotted lines mark the means, blue shaded region marks the extent of the reward zone. Note that cells that fire within 50 cm of one reward but not the other may have a mean relative distance of <0.698 radians (<50 cm); this is visible as the small fractions between the magenta lines marking ±50 cm from the start of the reward zone in (J, bottom) and (L, bottom). In (M), note that the fraction of cells above the shuffle shows a non-significant increase across task experience. **N)** Illustration of cross-correlation criterion to identify reward-relative remapping cells (top row) vs. track-relative cells (middle row) and non-reward-relative (non-RR) remapping cells (bottom row). Each example cell is also shown in (A), top row. Left of each cell: cross-correlation (xcorr) between trial-averaged spatial firing with reward zones circularly aligned, before vs. after the switch. Gray lines mark the mean (solid) and 95% confidence interval (dashed) of the shuffle for that cell. The offset of the xcorr maximum above the shuffle and the distance between relative spatial peaks in radians is listed next to each cell. Note that the reward-relative cell is maximally cross-correlated very close to zero lag, while the track-relative cell is anticorrelated at 0 lag in reward-aligned coordinates. Middle of each cell: spatial activity with reward zones aligned as in Fig. 2B; blue line marks the switch between reward zones. Right of each cell: trial-by-trial correlation matrix computed on the reward-aligned spatial activity. Note the more uniform structure of the correlation matrix for the reward-relative cell vs. the block-like matrices for the track-relative and non-RR remapping cell, which is the opposite of how these correlation matrices appear in original linear track coordinates (A, top row). **O)** Left: Distributions of Pearson correlation coefficients for each subpopulation between the cell’s trial-averaged activity maps pre- vs. post-switch in reward-aligned coordinates. The reward-relative (RR) population exhibits significantly higher correlations in these coordinates than the track-relative (TR) and non-RR remapping populations: RR vs. TR U = 67.7, p << 1e-12, Wilcoxon rank-sum test (n = 3104 RR cells, 3901 TR cells across 7 mice, 7 switch days); RR vs. non-RR remapping U = 53.8, p << 1e-12, Wilcoxon rank-sum test (n = 3104 RR cells, 2518 TR cells across 7 mice, 7 switch days). Right: Distributions of maximum xcorr offsets above the shuffle for each subpopulation, with the RR distribution thresholded at ±5 bins. **P)** Same as (O), but computed between trial-averaged activity of each subpopulation in original linear track coordinates. The TR population exhibits significantly higher correlations in these coordinates than the RR and non-RR remapping populations: TR vs. RR U = 69.1, p << 1e-12, Wilcoxon rank-sum test; TR vs. non-RR remapping U = 21.0, p << 1e-12, Wilcoxon rank-sum test.

**Figure S3.**
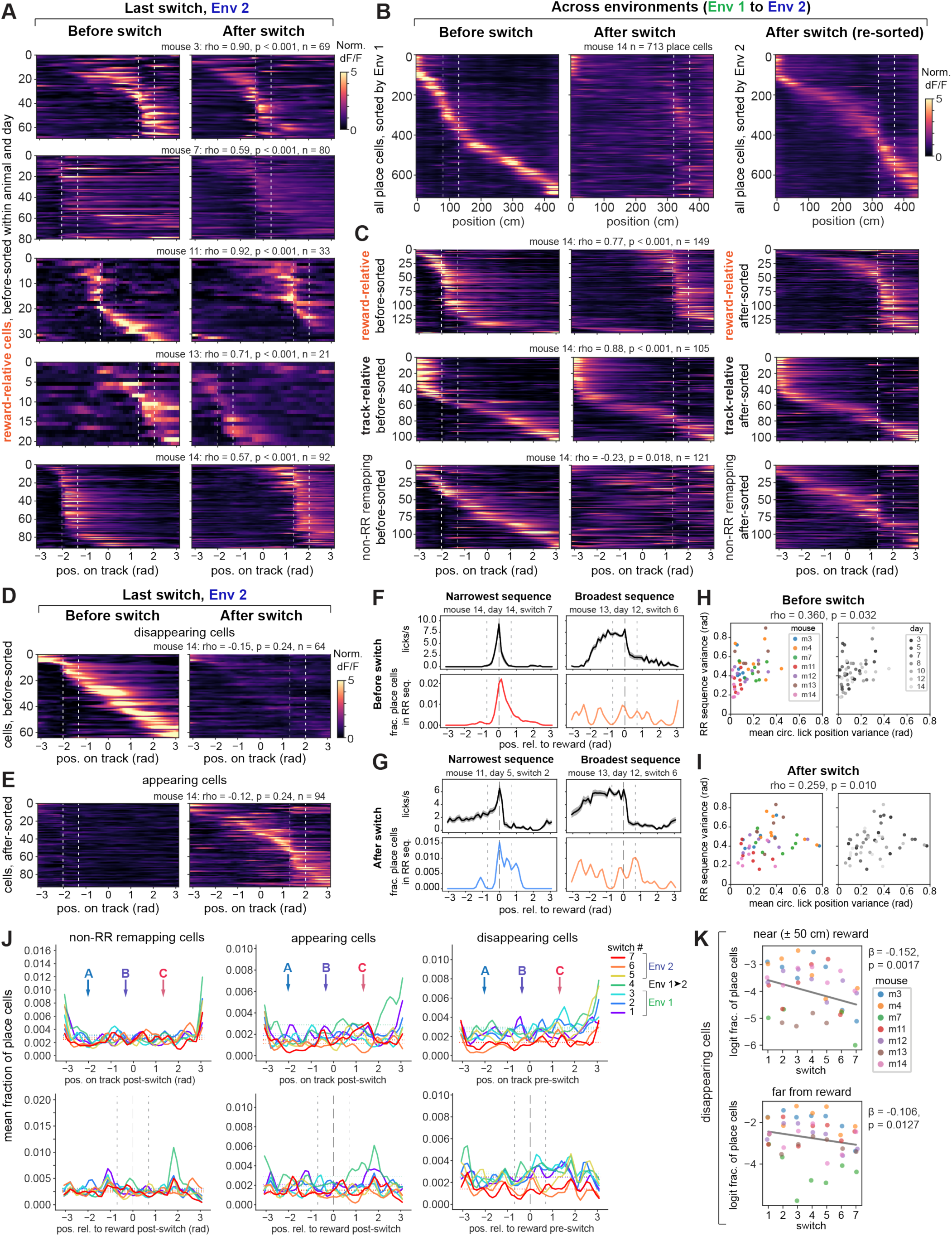
Sequences across animals, remapping types, and environments, related to Figure 3. A) Reward-relative sequences on the last switch (day 14) for the 5 mice not shown in Fig. 3A (organized as in Fig. 3A). B) Population level remapping upon the switch from Env 1 to novel Env 2 (day 8). All place cells identified in mouse m14 are shown, sorted in the left two panels by their order in Env 1 before the switch, and by their order in Env 2 in the right-most panel. Note population-level overrepresentation in both sorts as the bend away from the diagonal around the start of the reward zone. C) Remapping (or lack thereof) of specific subpopulations on the switch from Env 1 to Env 2 (day 8), shown for mouse m14. Top: reward-relative (RR) sequences are preserved across environments; Middle: a subset of track- relative place cells also retain their firing positions across environments, especially at the beginning of the track; Bottom: non-reward-relative, non-track-relative cells (“non-RR remapping”) with significant spatial information in both Env 1 and Env 2 instead remap globally, mirroring the overall population. The cells in the left two columns in (C) are sorted by their position in Env 1; the right-most column is sorted (and cross-validated) by position in Env 2, which improves the sort for the non-RR remapping cells but not the reward-relative or track-relative cells. D) Example sequences of disappearing cells from mouse m14. In (D) and (E), cross-validated sort was performed on the trials in which cells had significant spatial information (before for disappearing, after for appearing). E) Example sequences of appearing cells from mouse m14. F) Example licking (top) and distribution of reward-relative (RR) sequence peak firing positions during trials before the switch, aligned to the start of the reward zone. Left column: animal and day with the narrowest RR sequence (lowest circular variance); Right column: animal and day with the broadest sequence (highest circular variance). Sequence colors correspond to switch index as shown in (J). Note more distributed, less precise licking coinciding with a more distributed sequence that is less dense around the reward zone (right). Vertical gray dashed lines indicate the reward zone start and surrounding ±50 cm. G) Same as (F), but showing the narrowest and broadest sequences and corresponding licking for trials after the switch. **H-I)** Circular variance in licking positions is significantly correlated with RR sequence variance both before (H) and after the reward switch (I). Each dot is a session, colored by mouse (left; n=7 mice) or by switch day (right; n=7 days). Note no apparent trend over days. **J)** Mean distributions of peak firing positions across days (n=7 mice, s.e.m. omitted for clarity) for non-RR remapping cells (left), appearing cells (middle), and disappearing cells (right). Top row: sequences according to original linear track position, converted to radians. Bottom row: sequences according to position relative to the starts of reward zones. Horizontal dotted lines indicate the expected uniform distribution for each day. Vertical gray dashed lines indicate the reward zone start and surrounding ±50 cm. **K)** Both near (top) and far from (bottom) the start of the reward zone, the fraction of disappearing place cells out of total place cells decreases across experience. Fractions are shown as a logit transform here to account for values close to zero, and sessions with zero disappearing cells on any given day have been removed. Regression coefficients β and p-values are from linear mixed effects models with switch index as the fixed effect and mice as random effects. Non-RR remapping and appearing fractions did not show significant changes over experience (data not shown).

**Figure S4.**
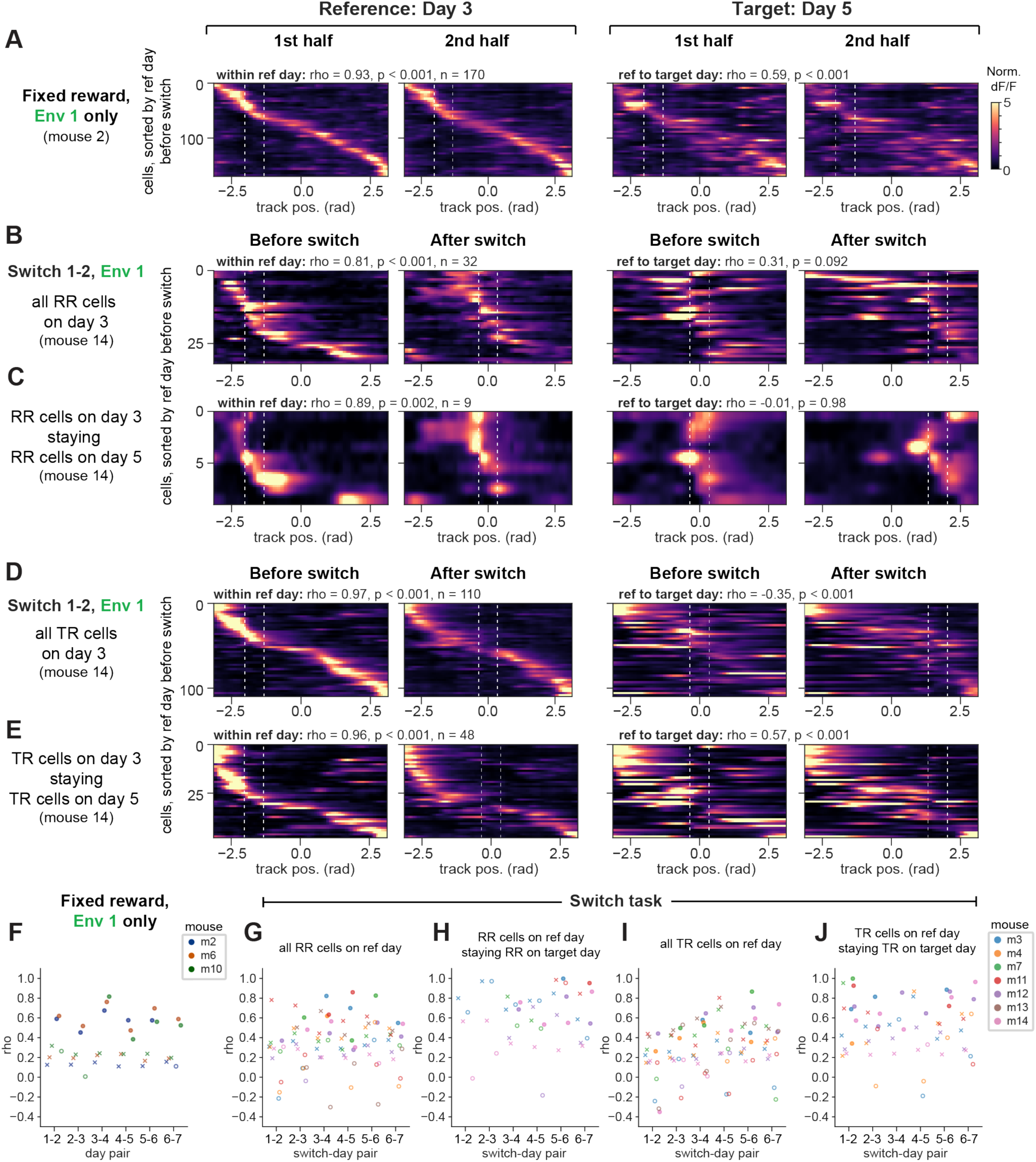
Drift in sequences tracked across days, related to Figure 4. **A)** An example place cell sequence tracked from days 3 to 5 in an animal (m2) from the fixed-condition cohort, which experienced the same reward zone and environment for all 14 days. Neurons in this animal were identified using the same criteria as reward-relative cells (i.e. their field position relative to reward had to be stable in terms of its peak position and spatial cross-correlation, from the 1st to the 2nd half of trials), but without a reward switch to identify reward-relative cells, this likely includes other cell categories. In (A-E), cells within an animal are sorted by their cross-validated peak activity in the 1st trial set (1st half, or before the switch) on the reference (ref) day, which is defined as the first day in each day pair. This sort then is applied to activity in the 2nd trial set on the reference day, as well as the 1st and 2nd trial sets on the target day (the second day in the pair). Also in (A-E), the sequence correlation coefficient, p-value relative to shuffle (two-sided permutation test), and n of cells tracked across the day pair are shown above each plot (within ref day correlation on the left, across-day correlation on the right; the across-day correlation is computed between the 2nd trial set on the ref day and the 1st trial set on the target day). In (A), note that drift is apparent across days in this fixed-condition animal as denoted by the drop in correlation coefficient, despite the cell ordering remaining significantly correlated across days. **B)** A reward-relative sequence identified in animal m14 on day 3 (switch 1) and tracked to day 5 (switch 2), agnostic of the cell categories for the cells on day 5. **C)** Reward-relative cells from day 3, m14 that were tracked to day 5 and remained reward-relative on day 5. **D)** A track-relative sequence identified in animal m14 on day 3 (switch 1) and tracked to day 5 (switch 2), agnostic of the cell categories for the cells on day 5. **E)** Track-relative cells from day 3, m14 that were tracked to day 5 and remained track-relative on day 5. **F)** Across-day circular-circular correlation coefficients for place cell sequences tracked across pairs of days in the fixed-condition cohort (n = 3 mice; 95 ± 54 tracked cells per day pair, mean ± s.d. across mice and day pairs). Place cells included for these sequences were identified as described in (A). In (F-J), coefficients are plotted with the upper 95^th^ percentile of 1000 shuffles of cell IDs (marked as x, jittered to the left of each set of coefficients, and color-coded by animal). Closed circles indicate coefficients with p < 0.05 compared to shuffle using a two-sided permutation test; open circles indicate p ≥ 0.05. **G)** Across-day circular-circular correlation coefficients for all cells in reward-relative sequences on each reference day tracked to each target day, for the animals performing the hidden reward zone switch task (n = 7 mice; 26 ± 20 cells, mean ± s.d. across mice and day pairs). **H)** Across-day circular-circular correlation coefficients for reward-relative cells on each reference day that were tracked and remained reward-relative on each target day (n = 7 mice; 8 ± 10 cells, mean ± s.d. across mice and day pairs). Note slightly higher correlation coefficients for this subset compared to (G), though the small numbers of cells that were both successfully tracked and remained reward-relative preclude a robust comparison. **I)** Across-day circular-circular correlation coefficients for all cells in track-relative sequences on each reference day tracked to each target day (n = 7 mice; 45 ± 37 cells, mean ± s.d. across mice and day pairs). **J)** Across-day circular-circular correlation coefficients for track-relative cells on each reference day that were tracked and remained track-relative on each target day (n = 7 mice; 15 ± 17 cells, mean ± s.d. across mice and day pairs).

**Figure S5.**
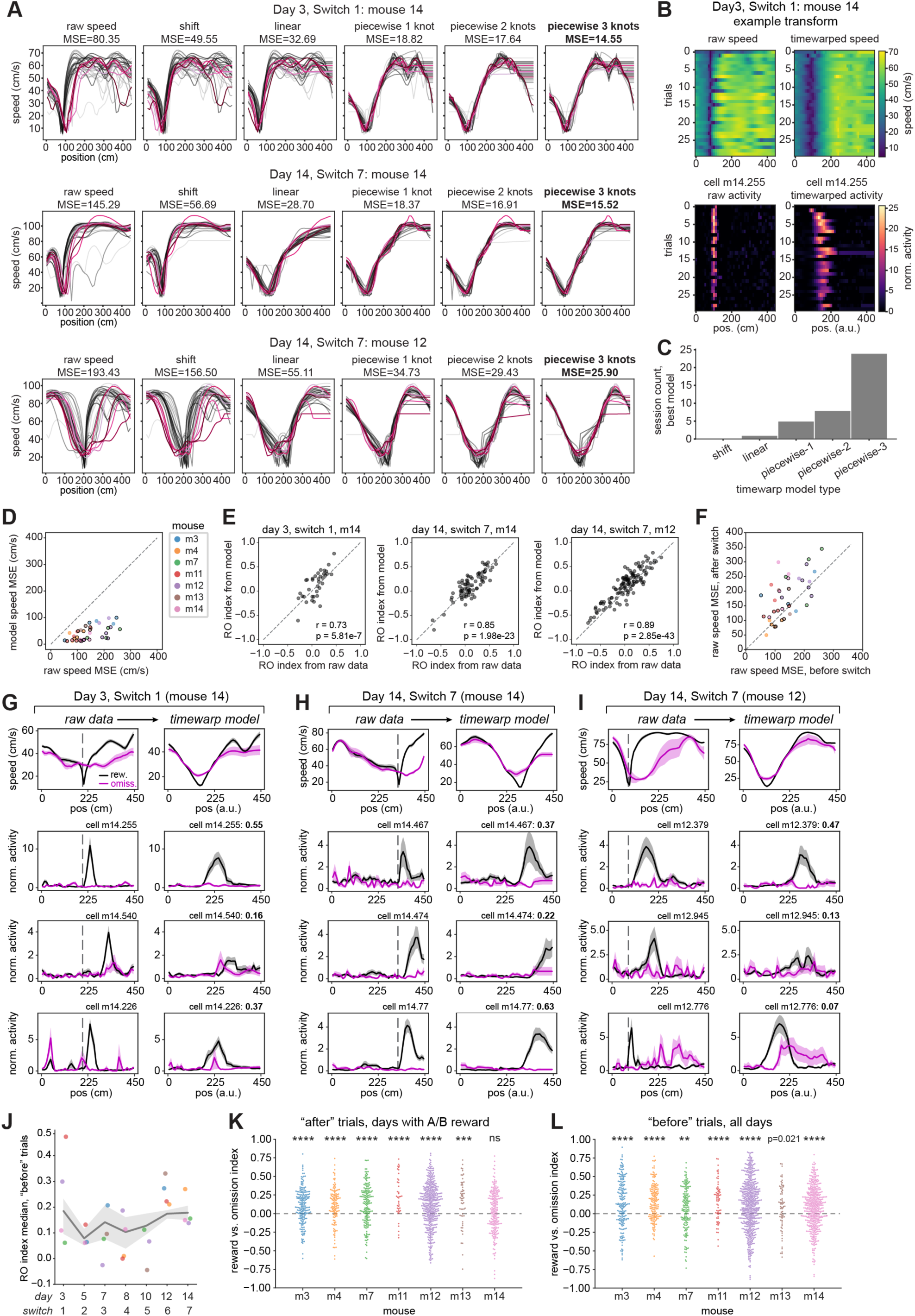
Implementation of time warp models and controls for analysis of rewarded vs. omission trials, related to Figure 5. **A)** Illustration of time warp model fitting procedure to running speed profiles. Shades of black indicate speed on rewarded trials, shades of magenta indicate omission trials. Left to right: raw speed, followed by the transformed speed from fitting 5 model types to identify a model for best fit, as assessed by mean squared error (MSE) across the transformed trial-by-trial speed profiles. Bold text indicates the best-performing model. Top to bottom: the same example sessions shown in Fig. 5A-C: first reward switch on day 3, mouse 14; last reward switch on day 14, mouse 14; last reward switch on day 14, mouse 12. **B)** Example transformation using the piecewise 3 knots model fit on the speed from day 3, mouse 14 (shown in A, top). Top left: raw trial-by-trial speed. Top right: time warp transformed speed. Bottom left: raw, binned deconvolved activity (normalized to the mean of the session) for example cell m14.255, which is also shown in Fig. 5A (top row of cells). Bottom right: time warp transformed neural activity of cell m14.255. **C)** Frequency with which each model type was selected as maximally aligning speed according to MSE. For further analysis, we applied piecewise-3 to all data as this was the most common best model, to avoid variance in results due to model type for individual sessions. **D)** Time-warp model fitting significantly reduces MSE across trial-by-trial speed profiles compared to the original speed data (model aligned speed vs. raw speed: W = 0, p = 7.74e-8, Wilcoxon signed-rank test, n = 38 sessions across 7 mice). Each dot is a session, colored by mouse. Black edges indicate sessions with the reward zone at location “A” or “B”, included for the analysis shown in Fig. 5. In (D-F), gray dashed line marks the unity line. **E)** Reward vs. omission index (RO index) calculated from the model-transformed neural data, shown for the same sessions in Fig. 5D, is highly correlated with the RO index calculated from the original neural data (Pearson correlations). This indicates that the time warp model scales and aligns the firing rate curves between rewarded and omission trials but does not distort their relationship. **F)** MSE between trial-by-trial raw speed profiles before vs. after the reward switch (within each session with at least 3 omission trials before the switch). Each dot is a session, colored by animal as in (D). Black edges indicate sessions included for analysis in Fig. 5. Speed MSE after the switch is significantly higher than before the switch (thus rewards and omissions are more difficult to compare on these trials): W = 130, p = 4.87e-4, Wilcoxon signed-rank test, n = 38 sessions across 7 mice. **G)** Raw and model-aligned speed and neural activity of the same session and example cells shown in Fig. 5A, but for the trials after the reward switch (“after” trials). RO indices shown in bold text (top, far right of right column panel) are calculated from the “after” trials. Note preference of cells to fire more following rewards versus omissions, with the caveat that the speed profiles are more different than during the “before” trials. **H)** Same session and example cells as Fig. 5B, but for the “after” trials. **I)** Same session and example cells as Fig. 5C, but for the “after” trials. **J)** A linear mixed effects model shows no significant effect of day (β= 0.01, p = 0.73), original MSE between mean speed on rewarded and omission trials (β= -0.001, p = 0.49), MSE of the time warp model fit (β = 0.002, p = 0.86), or the interactions of these terms (fixed effects) on the median RO index (random effects are mouse identity). Each dot shows the median RO index of the sessions included in Fig. 5 for each mouse, colored by mice as in (D). Gray line shows the model prediction ± 95% confidence interval. **K)** Reward vs. omission index across all switch days (sessions) where the reward zone was at “A” or “B”, on the trials after the switch, for the population of reward-relative cells that have spatial firing peaks following the start of the reward zone within each mouse (n = 7 mice). In (K-L), significance level is set at p<0.007 with Bonferroni correction. **p<0.007, ***p<0.001, ****p<0.0001, one-sample Wilcoxon signed-rank test against a 0 median. m3: median = 0.10, W = 5.01e3, p = 3.31e-9, n = 5 days, 197 cells; m4: median = 0.13, W = 2.10e3, p = 3.70e-6, n = 4 days, 126 cells; m7: median = 0.14, W = 4.21e3, p = 7.58e-8, n = 5 days, 177 cells; m11: median = 0.22, W = 106, p = 2.61e-5, n = 3 days, 41 cells; m12: median = 0.13, W = 2.57e4, p = 1.55e-13, n = 5 days, 419 cells; m13: median = 0.21, W = 377, p = 2.12e-4, n = 5 days, 58 cells; m14: median = 0.04, W = 9.84e3, p = 0.085, n = 3 days, 213 cells. **L)** Reward vs. omission index across all switch days (sessions) with at least 3 omission trials, at any reward zone before the switch, within each mouse (complementary to Fig. 5E, n = 7 mice). m3: median = 0.13, W = 4.99e3, p = 4.86e-8, n = 5 days, 191 cells; m4: median = 0.15, W = 2.63e3, p = 7.11e-12, n = 5 days, 165 cells; m7: median = 0.09, W = 4.44e3, p = 2.08e-3, n = 4 days, 157 cells; m11: median = 0.20, W = 418, p = 3.96e-6, n = 6 days, 68 cells; m12: median = 0.13, W = 4.66e4, p = 1.96e-13, n = 7 days, 541 cells; m13: median = 0.10, W = 902, p = 0.021, n = 5 days, 72 cells; m14: median = 0.09, W = 3.35e4, p = 9.13e-9, n = 6 days, 442 cells.

**Figure S6.**
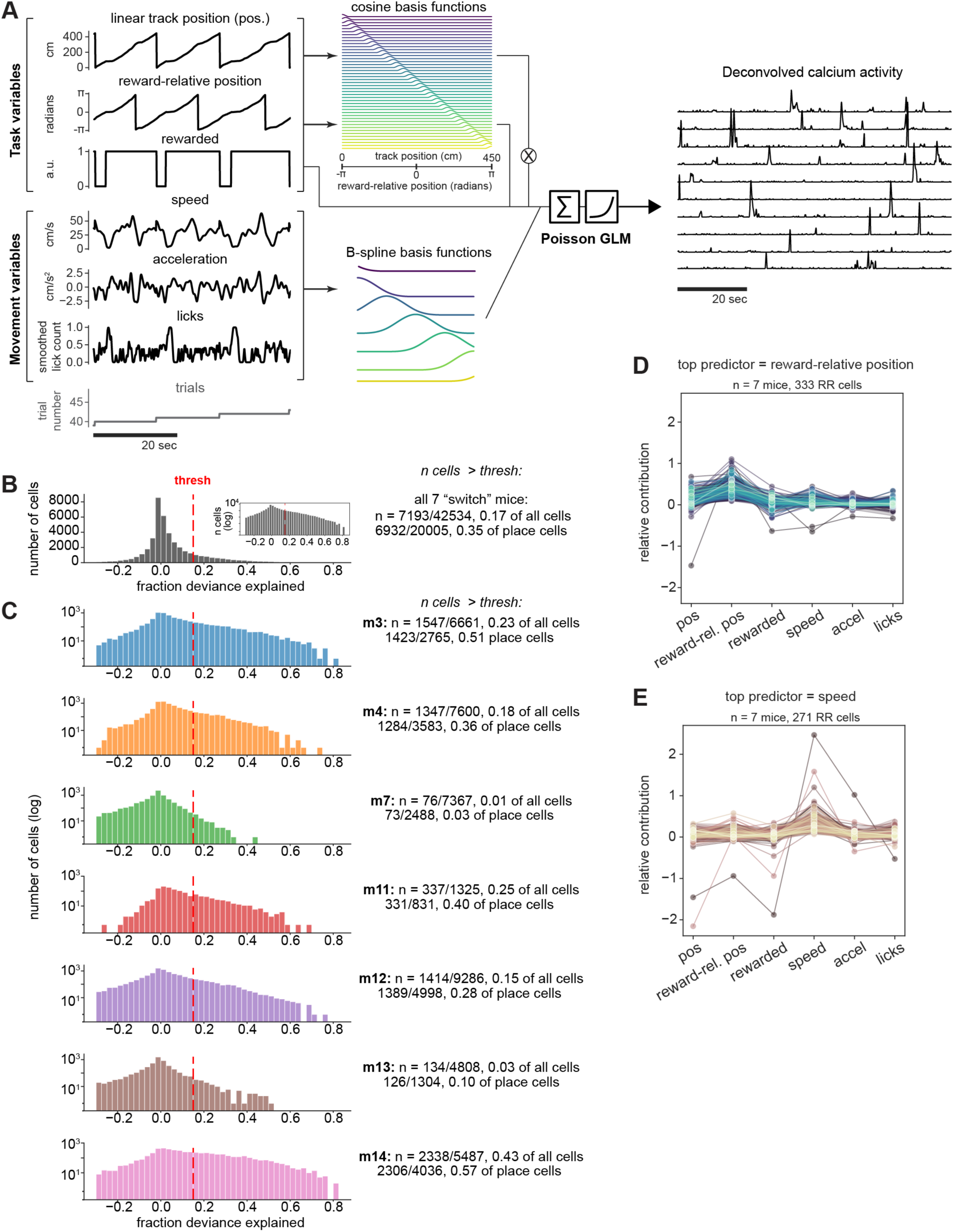
Generalized linear model (GLM) implementation and additional quantification, related to Figure 6. **A)** Schematic of the Poisson GLM to predict deconvolved calcium activity from task and movement variables. Linear track position and reward-relative position are both transformed into cosine basis functions tiling the space in each set of coordinates (45 cosine bumps for the 45 10-cm spatial bins used). A binary representing whether reward was received on each trial (“rewarded”) is multiplied with the linear track position basis functions to represent the interaction between reward received and position. Speed, acceleration, and smoothed lick count are quantile-transformed into B-splines with 5 knots and 3 polynomial degrees (providing 7 splines total) which smoothly tile the range of speeds, accelerations, and licking within each animal. Trial identities are not provided as an input but are used to group data for the training, cross-validation, and test sets (see Methods). **B)** Quantification of model performance (fraction deviance explained on test data) across all animals and cells. Inset shows the histogram on a logarithmic scale. Red dashed line indicates the fit threshold of 0.15 fraction deviance explained used to select neurons for further analysis. Number and fraction of cells that exceeded the threshold is listed relative to all cells and relative to place cells, across all switch days. **C)** Quantification of model performance within each animal (n = 7 mice), shown on a logarithmic scale. **D)** Relative contributions of each GLM variable within reward-relative cells which were maximally predicted by reward-relative position. Each line and set of dots is an individual cell. **E)** Relative contributions of each GLM variable within reward-relative cells which were maximally predicted by running speed, shown as in (D).

## Notes

### Competing Interest Statement

The authors have declared no competing interest.

### Summary of Updates

Two main figures and two supplemental figures have been added with additional results.

